# A meta-analysis of rice phosphoproteomics data to understand variation in cell signalling across the rice pan-genome

**DOI:** 10.1101/2023.11.17.567512

**Authors:** Kerry A Ramsbottom, Ananth Prakash, Yasset Perez Riverol, Oscar Martin Camacho, Zhi Sun, Deepti J. Kundu, Emily Bowler-Barnett, Maria Martin, Jun Fan, Dmytro Chebotarov, Kenneth L McNally, Eric W Deutsch, Juan Antonio Vizcaíno, Andrew R Jones

## Abstract

Phosphorylation is the most studied post-translational modification, and has multiple biological functions. In this study, we have re-analysed publicly available mass spectrometry proteomics datasets enriched for phosphopeptides from Asian rice (*Oryza sativa*). In total we identified 15,522 phosphosites on serine, threonine and tyrosine residues on rice proteins.

We identified sequence motifs for phosphosites, and link motifs to enrichment of different biological processes, indicating different downstream regulation likely caused by different kinase groups. We cross-referenced phosphosites against the rice 3,000 genomes, to identify single amino acid variations (SAAVs) within or proximal to phosphosites that could cause loss of a site in a given rice variety. The data was clustered to identify groups of sites with similar patterns across rice family groups, for example those highly conserved in Japonica, but mostly absent in Aus type rice varieties - known to have different responses to drought. These resources can assist rice researchers to discover alleles with significantly different functional effects across rice varieties.

The data has been loaded into UniProt Knowledge-Base - enabling researchers to visualise sites alongside other data on rice proteins e.g. structural models from AlphaFold2, PeptideAtlas and the PRIDE database - enabling visualisation of source evidence, including scores and supporting mass spectra.

## Introduction

Rice is one of the most important crops for human nutrition, acting as staple food for around a third of the global human population [1]. Asian domesticated rice, *Oryza sativa*, has historically been sub-categorised into two major varietal groups: Japonica and Indica, although further sub-divisions have also been proposed, including Aus and Admixed families. There is great genetic diversity both within and between varietal groups. Major efforts are underway to understand that diversity through genomic techniques, and to exploit diversity to find alleles conferring desirable traits (such as resistance to biotic and abiotic stresses), which could be bred into high yielding varieties. The genome sequence of a reference Japonica variety, Nipponbare, was sequenced by the International Rice Genome Sequencing Project (IRGSP), with a first release of gene models in 2005 [2]. A group led from China also sequenced a reference Indica variety (IR-93), and independently annotated gene models [3]. Despite Indica rice varieties accounting for around six times the size of international market as Japonica rice varieties [4], the Nipponbare assembly is generally considered the “canonical reference” genome for research and breeding efforts.

There are two current, non-synchronised annotations of the *Oryza sativa* Japonica (Nipponbare variety) genome assembly: the Rice Genome Annotation Project at Michigan State University (MSU) [5] and the Rice Annotation Project Database (RAP-DB) [6]. MSU gene models are no longer updated but still used frequently in research projects and cited in publications. RAP-DB is regularly updated, and serves as the source for gene models loaded into other databases such as Ensembl Plants and Gramene [7, 8] (the two databases being mostly synchronised and using the same underlying technologies), and the source protein sequences for UniProtKB (UniProt Knowledge-base) [9], the most popular protein knowledge-base. Over several years UniProtKB has performed some manual curation, where improvements can be identified in protein sequences, meaning that UniProtKB protein sequences are not identical to Ensembl Plants/Gramene. Other key initiatives and datasets include the rice 3,000 genomes project [10], which provides a resource for understanding genetic variants within *Oryza sativa*. More recently, new rice “platinum standard” genomes are being sequenced sequenced [11, 12], with new predicted gene models for these varieties now available in Ensembl Plants and Gramene.

Genetic variation data and GWAS analyses can be key for identifying candidate genes or chromosomal regions associated with traits of interest. However, discovery of a SNP (Single Nucleotide Polymorphism) significantly associated with a trait can give only limited information about associated biological function or mechanism. For example, to understand why a given trait confers stress resistance involves understanding the function of proteins, and the pathways and networks they are involved with. A key component relates to understanding cell signalling, such as fast responses to the detection of stress, via reversible post-translational modifications (PTMs) of proteins. The most widely studied reversible modifications include phosphorylation (by far the most studied one, and our primary focus here), acetylation, methylation, and attachment of small proteins, such as ubiquitin and SUMO. There is increasing evidence that sites of PTMs can be important alleles for breeding efforts, examples including the “green revolution” DELLA genes that have an altered response to the gibberellin hormone, via loss of PTM sites [13] and root branching towards water controlled by SUMOylation [14].

In this work, we aim to provide a high-quality resource providing phosphorylation sites in rice. Phosphosites on proteins are detected and localised on a large scale using tandem mass spectrometry (MS), via “phosphoproteomics” methods. These methods generally involve the proteins extracted from samples being digested by enzymes such as trypsin and phosphorylated peptides being enriched in these samples using reagents such as TiO2, or other metal ions, attached to a column (affinity chromatography), to which phosphate binds preferentially. These bound peptides are then eluted and analysed using liquid chromatography-mass spectrometry (LC-MS/MS) [15]. The tandem MS data is then usually queried against a protein sequence database, via a search algorithm. Scores or statistics are calculated for the confidence that the correct peptide sequence has been identified (including the mass of any PTMs), and then for PTM-enriched data, a second step is usually performed to assess the confidence that the site of modification has been correctly identified, if there are multiple alternative potential residues in the peptide. We have recently published an approach to assess the global false localization rate (FLR) of PTMs using searches for PTMs on decoy amino acids, and demonstrated its importance for controlling multiple sources of error in the analytical pipeline [16]. We have also extended the model to demonstrate how to combine evidence coming from multiple spectra, and to combine evidence and control FLR across multiple studies[17].

Many papers that report phospho-proteomes do not adequately control for site FDR, and just use *ad hoc* score thresholds for peptide identification or site localisation scores. For example, in the popular PhosphoSitePlus resource we estimated that around ∼67% of the phosphosites reported in the database are likely to be false positives, and those with only an observation from one single study are very unlikely to be true [18]. To provide FDR-controlled data on phosphosites to the research community requires reprocessing MS data, using well controlled statistical procedures, and applying *post hoc* approaches to control FDR when aggregating data across multiple studies.

There are several online databases for gaining information about PTMs in plants, sourced from published studies. The Plant PTM Viewer [19] aggregates results from published studies that used PTM enrichment and MS, and has good coverage of studies for a range of PTM types, including phosphorylation, acetylation, ubiquitination and several others. Over 13,000 rice proteins are reported to be modified, and >27,000 *Arabidopsis thaliana* proteins, mapped to ∼15,000 genes (translated from 54,000 transcripts of the 27,600 coding genes). For detailed study of the *A. thaliana* phosphoproteome, there also exists the PhosPhAT database [20], which similarly loads phosphoproteomics data from published studies on *Arabidopsis*, containing evidence for 55,000 phosphorylation sites on ∼9,000 *A. thaliana* proteins. Plant PTM Viewer and PhosPhAT are useful for community resources, although by loading data from published studies (rather than re-processing data), are likely to contain variable data quality and cannot control for FLR across multiple datasets.

UniProtKB is a leading cross-species resource for studying protein function, including extensive expert manual curation. For PTM-related data, UniProtKB mostly loads data by curating individual studies, and has not previously loaded large-scale MS data reporting on plant PTMs. PTMs are reported on just 320 rice proteins in the UniProtKB and on 2,763 *Arabidopsis* proteins (October 2023, Release 2023_04).

The PRIDE database at the European Bioinformatics Institute (EMBL-EBI) is the largest MS-based proteomics data repository [21]. PRIDE is leading the ProteomeXchange (PX) consortium of proteomics resources, whose mission is to standardise open data practices in proteomics worldwide [22]. PeptideAtlas is also a PX member, focused on the consistent reanalysis of datasets [23] for a variety of species, including recent builds for Arabidopsis [24]. Widespread public deposition of proteomics data in PX resource now enables meta-analysis studies to be performed, by reanalysing groups of related datasets. As part of the “PTMeXchange” project, our consortium aims to complete a large-scale re-analysis of public PTM enriched datasets, using robust analysis pipelines incorporating strict FDR control, and correction for FDR inflation in meta-analyses.

In this work, we have re-analysed phospho-enriched rice datasets and integrated results into PeptideAtlas, PRIDE and UniProtKB, for visualisation of the confident phosphosites alongside other data on rice proteins. Downstream analysis is also performed on the confident sites to identify PTMs which may be of biological interest. These analyses include investigations on common motifs seen around the phosphosites, pathway enrichment analysis for these motifs and analysis of single amino acid variations (SAAVs) identified close by to the confident phosphosites. From these analyses, we aim to identify rice phosphoproteins which may be of biological interest.

## Methods

### Phosphosite identification

The ProteomeXchange Consortium [25] was used to identify suitable rice phosphoproteomics datasets, via the PRIDE repository [26]. From this, 111 proteomics datasets were identified for the *Oryza sativa* species. Of these, 13 were identified as being enriched in phosphopeptides and then potentially suitable for reanalysis: PXD000923 [27], PXD001168 [28], PXD000857 [28], PXD002222 [29], PXD001774 [30], PXD004939 [31], PXD005241 [32], PXD004705 [33], PXD002756 [34], PXD012764 [35], PXD007979 [36], PXD010565 [37] and PXD019291 [38] (Supplementary table 1). These 13 datasets were investigated further to evaluate their quality for use within the phosphoproteomics reanalysis. It was found that PXD001168, PXD001774 and PXD010565 contained very few phosphopeptides for the size of the dataset, these datasets were therefore excluded from the analysis. PXD007979 was identified to be a meta-analysis of the PXD002222 and PXD000923 datasets and was therefore also excluded. Finally, PXD000857 was excluded as this is a relatively old dataset containing only one raw file. This resulted in 8 high quality datasets being carried forward for the phosphopeptide re-analysis. Sample and experimental metadata were manually curated and adhering to the Sample-Data Relationship Format (SDRF)-Proteomics file format [39].

The search database was created consisting of protein sequences derived from the MSU Rice Genome Annotation Project, the Rice Annotation Project Database (RAP-DB), including both translated CDS and predicted sequences, and UniProtKB, including both reviewed and unreviewed sequences. A fasta file was generated from the combination of these databases, if a protein sequence occurs in more than one resource, RAP-DB was used as the primary identifier, this was then followed by any other IDs for that protein. cRAP contaminant sequences were also added to the database (https://www.thegpm.org/crap/, accessed April 2022) and decoys across all protein and contaminant sequences were generated for each entry using the de Brujin method (with k=2) [40]. The database was deposited in PRIDE along with the reprocessed data files (PRIDE ID: PXD046188).

The analysis was conducted using the pipeline as previously described [16]. Using the Trans-Proteomic Pipeline (TPP) [41, 42], the dataset files were first searched using Comet [43]. The resulting files were then combined and processed using PeptideProphet [44], iProphet [45], and PTMProphet [46], for each dataset. The files were searched with the variable modifications: Oxidation (MW), N-terminal acetylation, ammonia loss (QC), pyro-glu (E), deamination (NQ) and phosphorylation (STYA). Phosphorylation on alanine was included as a decoy to estimate false localisation rate (FLR), using the count of pAla identified, following the methods previously described by our group [16]. Carbamidomethylation (C) was used as a fixed modification and the iTRAX8plex label was included for the search on the PXD012764 dataset. Maximum missed cleavage used was 2, with a maximum number of modifications per peptide of 5. Table 1 outlines the datasets used and the tolerance parameters used for each dataset.

**Table 1:**
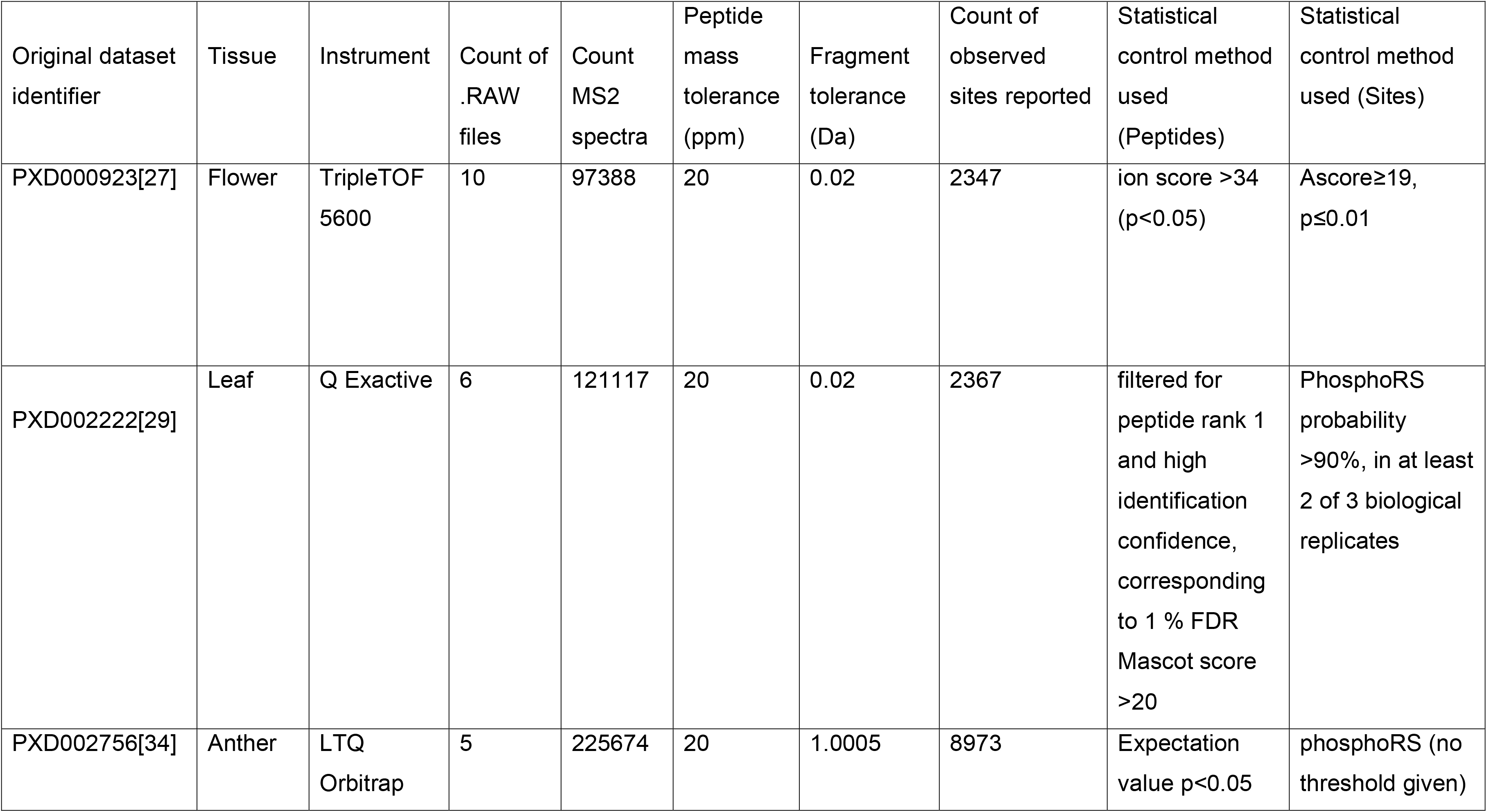
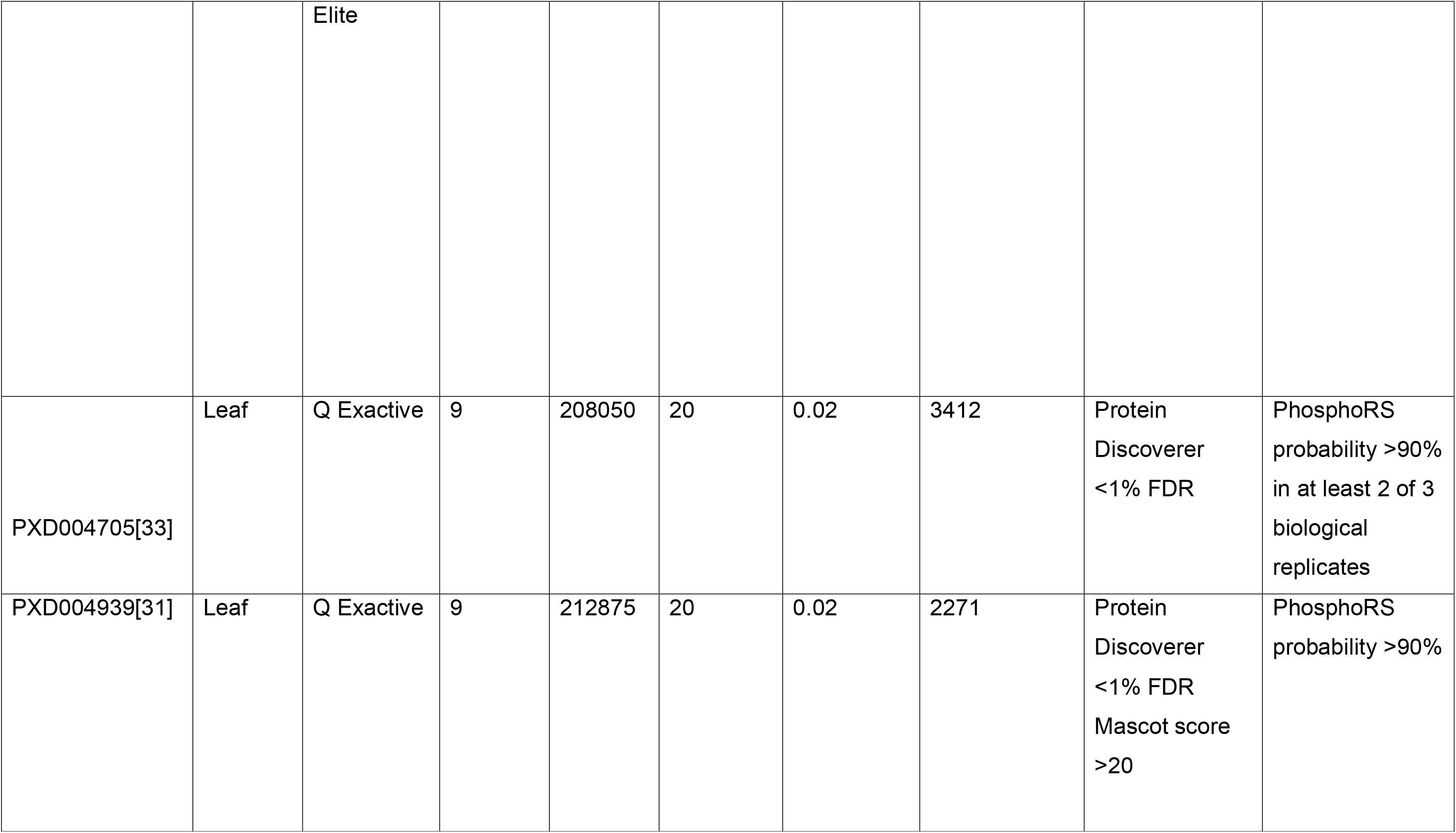
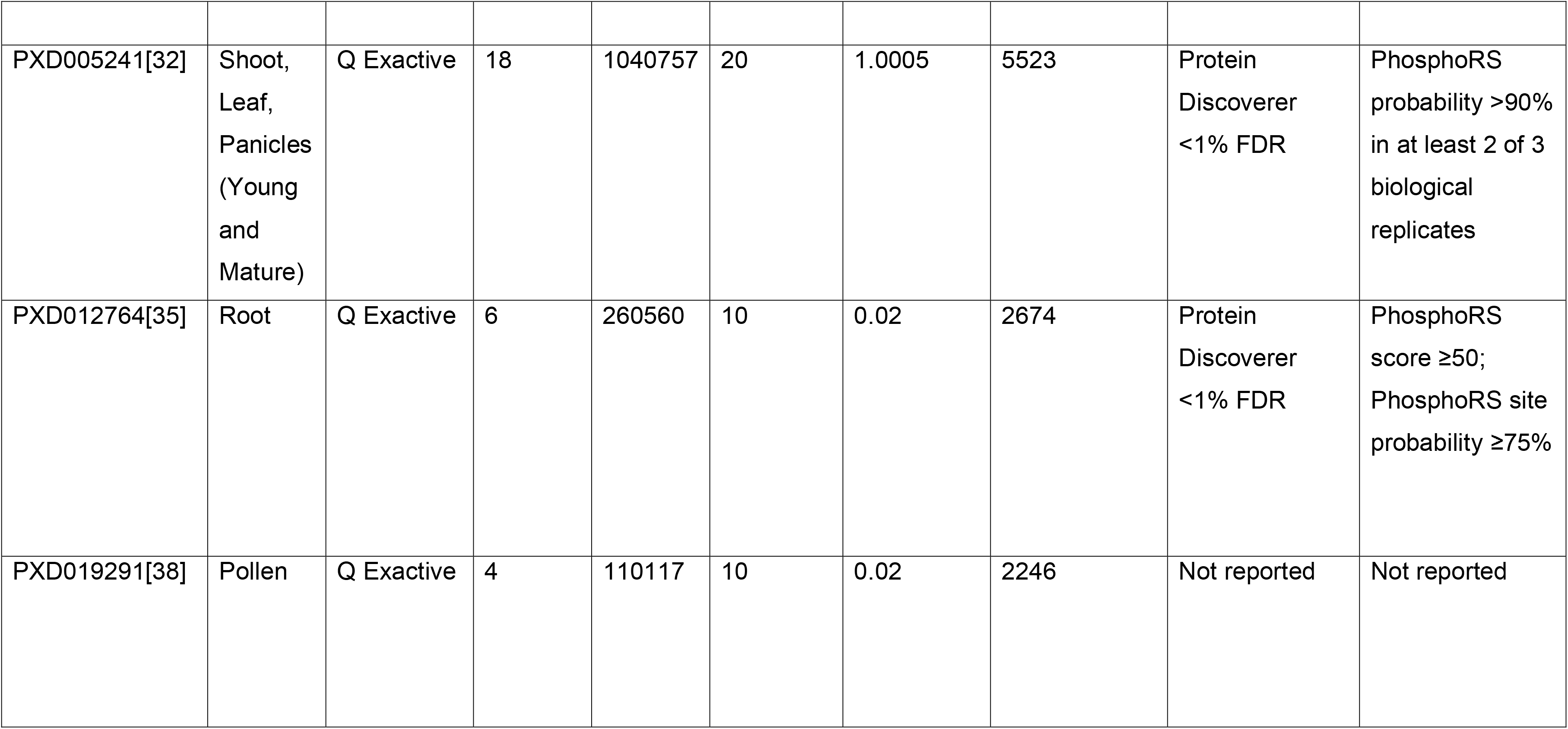
Tolerance parameters used for comet search for each dataset.

The data files obtained from searching with TPP were processed by custom Python scripts (https://github.com/PGB-LIV/mzidFLR). The data was analysed in the same way as in a previous study [16, 17]. The global FDR was calculated from the decoy counts and the peptide-spectrum matches (PSMs) were filtered for 1% PSM FDR. From these filtered PSMs, a site-based file was generated giving individual localisation scores for each phosphosite found on each PSM, removing PSMs not containing a phosphate, decoy PSMs and contaminant hits. These site-based PSMs were ordered by a *combined probability*, calculated by multiplying the PSM probability by the localisation probability.

It is common to observe many PSMs giving evidence for sites on the same peptidoform, where a peptidoform is a peptide sequence with a specific set of modified residues. In previous work [17], we have shown that collapsing results to the peptidoform-site level simply by taking the maximum final probability was sub-optimal as many of the high scoring decoy (and thus false) hits are supported by only a single PSM. We therefore applied a statistical model for multiple observations of a PTM site, using a binomial adjustment of the PTM probabilities to collapse these results by protein position [17]. This adjustment considered the number of times a specific site has been seen and the number of times this same site has been seen as a phosphosite, allowing us to give weight to those sites that are supported by multiple PSMs.

The global FLRs for all the datasets were estimated using the identification of phosphorylated Alanine (pAla) as a decoy. These are known to be false localisations and can therefore be used to estimate the FLR, following the method previously established [16], alongside the binomial adjustment. Global FLR was estimated for every ranked site, across all PSMs and in the collapsed protein position site-based format, from which we can later apply a threshold at the lowest scoring site that delivers a desired global FLR (e.g. 1%, 5% or 10%), similar to the q-value approach for standard proteomic database searching.

When aggregating data across multiple studies, we must control the inflation of FLR. FLR inflation is observed when the same correct sites are seen identified across multiple studies and tend to accumulate slowly, whereas each study reports different and random false positives which then accumulate rapidly as more datasets are added. PTM localisation has been shown to be incomparable between independent studies [17]. As a result, we developed an empirical approach to categorise sites based on the observations of sites across datasets at different thresholds; Gold, Silver and Bronze. Gold represents sites seen in *n* dataset with <1% FLR, Silver represents sites seen in *m* datasets with <1% FLR and Bronze represents any other sites passing <5% FLR. The values for *n* and *m* can be set empirically in a PTM “build” based on the number of datasets and the counts of decoys following the aggregation of multiple datasets, and application of possible values of n and m. As our rice build contains eight datasets, we categorised Gold sites as seen in more than one dataset with <1% FLR and Silver as only one dataset with <1% FLR. We then calculated the counts of pAla sites within these sets, allowing us to estimate the resulting FLR following dataset merging in the different categories.

### Dataset deposition and visualisation

The reprocessed data has been deposited in PRIDE in mzIdentML format [47], as well as SDRF-Proteomics files, tab-separated text formatted files (one per dataset) containing sites detected per PSM, and sites detected for each peptidoform, following the collapse processed described above (PRIDE ID: PXD046188). To load all phosphosites into UniProtKB, the identified peptides were mapped to the canonical protein sequences within the proteome (UP000059680) using an exact peptide sequence match, following theoretical tryptic digest. The phophosites can be viewed in the Protein APIs, in the Feature Viewer under Proteomics track and in the entry page. Decoy sites were not loaded to avoid misinterpretation.

All results have also been loaded as a PeptideAtlas build, available at https://peptideatlas.org/builds/rice/phospho/. The PeptideAtlas interface allows browsing of all modified peptides from these datasets, including those passing and not passing the above thresholds. All localization probabilities for all PSMs are displayed, along with links to the original spectra that may be visualized in the PeptideAtlas interface. The corresponding mass spectra in PeptideAtlas and PRIDE (https://www.ebi.ac.uk/pride/archive/usi) can be referenced and accessed via their Universal Spectrum Identifiers [48].

## Downstream analysis

### Motif and pathway enrichment analysis

All eight datasets were further investigated using motif and pathway enrichment analysis. Once the confident phosphosites (5% pAla FLR) from each of the datasets had been combined and given an FLR ranking (Gold, Silver, Bronze) the enriched motifs surrounding phosphosites were identified using the R [49] package *rmotifx* [50]. 15mer peptides were generated surrounding each of the identified phosphosites. These phosphopeptides were compared against a background of 15mer peptides with STY at the central position of the 15mer, and matched to the central residue of the phosphosite motif, to identify the enriched motifs seen around the confident phosphosites.

The proteins containing these enriched motifs were then carried forward for pathway enrichment analysis using ClusterProfiler [51]. The proteins containing each enriched motif was compared against all phosphoproteins in the search database. Similarly, a comparison was made between all phosphoproteins containing any enriched motif, for each of the FLR ranking categories, against the background of all phosphorylated proteins in the search database.

### SAAV Analysis

We also explored the phosphosites across all datasets we have re-analysed and how SAAVs may potentially affect these sites. We compiled a list of unique phosphosites (by sites on unique peptides) from the confident phosphosites (5% pAla FLR) across all eight searches and created a matrix showing which sites were seen in each dataset. These were then also mapped to the relevant protein sites in the three search databases: MSU, RAP-DB and UniProtKB. We mapped this list of unique phosphosites to known SAAV positions for the 3,000 rice varieties using the Rice SNP-Seek Database API (Application Programming Interface), for those sites that mapped to the MSU database. We categorised the phosphosites with relation to the SAAV sites; where “Category 1” = SAAV at the same position as a phosphosite, “Category 2” = SAAV at the +1 position to a phosphosite, “Category 3” = SAAV at the −1 position to a phosphosite and “Category 4” = SAAV at +/-5 amino acids from a phosphosite (and not in Category 1, 2 or 3). All other sites were assigned “Category 0”. For each phosphosite in the unique list across all datasets, the nearest SAAV to each phosphosite was identified and categorised. For those protein phosphosites with SAAV data available, we then investigated which alleles carried the SAAV and the minor allele frequencies for each site. We also added in annotation to show the genes involved, obtained using the Oryzabase database [52]. From this analysis, we could identify candidate sites of potential biological importance, which may be disrupted due to SAAVs.

A protein multiple sequence alignment was created using Clustalx 2.1, for the example protein Os09g0135400, versus the same locus in 15 other *Oryza sativa* genomes, which have been annotated and deposited in Ensembl Plants and Gramene [53, 54]. The association of gene models from different cultivars to be the same locus (a “pan gene cluster”) was created using the GET_PANGENES pipeline [55].

## Results and Discussion

### Phosphosite identification

First, we re-analysed each of the rice datasets identified as suitable for phosphosite identification using TPP as explained in Methods (Figure 1). Our scripts were used to calculate the FLR across confident PSMs identified (filtered for 1% FDR). We next collapsed the sites by protein position and remove duplicated hits. The FLR estimation was then recalculated on these collapsed sites, ordering by the calculated probability that a site had been observed, considering evidence across multiple PSMs (see Methods). The counts of phosphosites passing at different FLR thresholds are shown in Table 2 and Figure 1a (counts are derived from the unique combinations of a peptidoform and phosphosites within those peptidoforms, i.e. not accounting for some peptidoforms mapping to more than one genomic locus (gene)).

**Figure 1:**
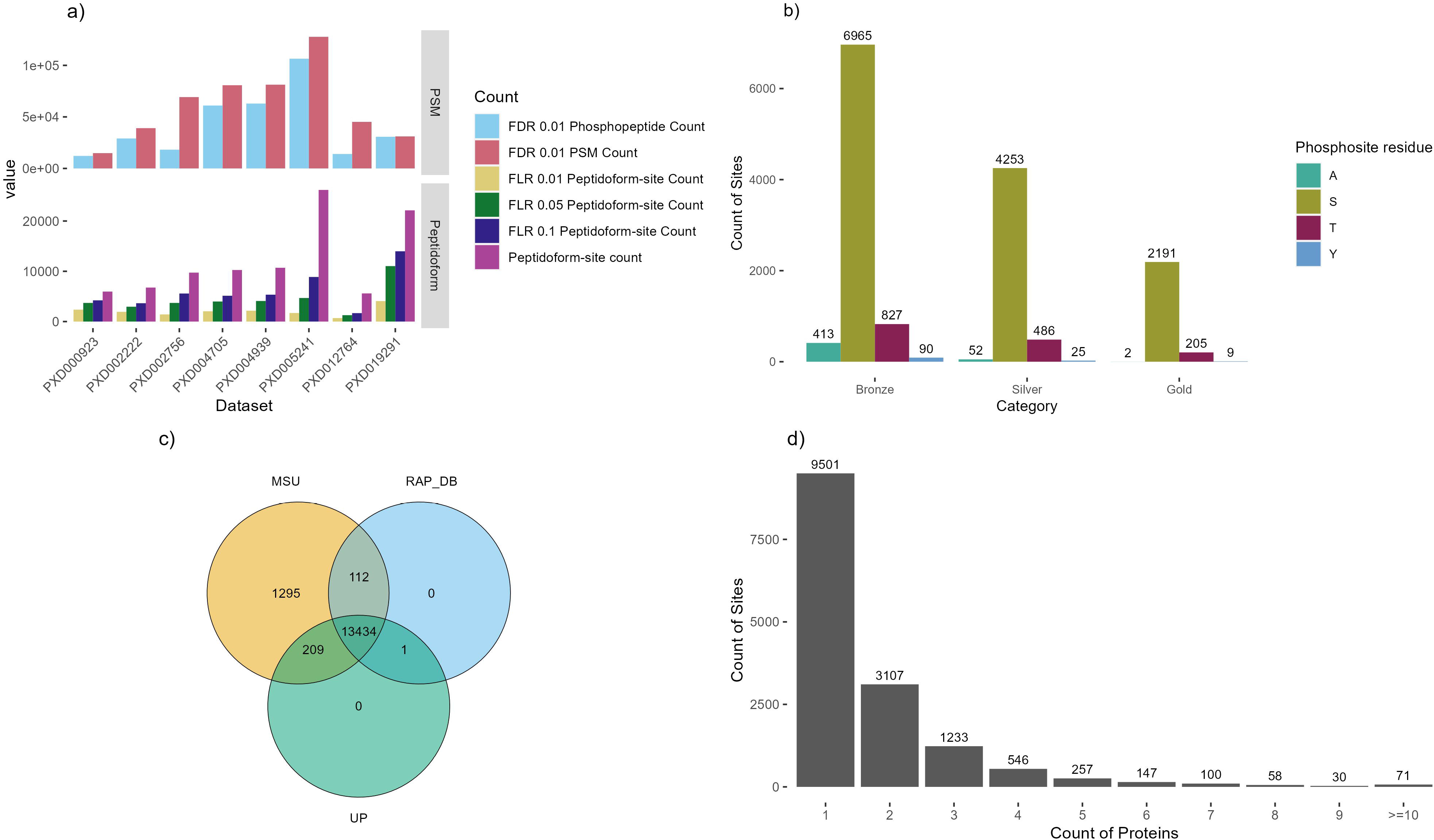
a) Counts of phosphosites before and after (peptidoform) collapse for removing redundancy at three FLR levels per dataset; b) The count of phosphosites in each of the three categories (Gold-Silver-Bronze) per residue where A is the decoy Alanine; c) The overlap of phosphosites reported per protein database: MSU, RAP-DB or UniProtKB (UP); d) Counts of phosphosites observed across different protein counts.

**Table 2:** PSM counts for each dataset at 1% FDR, Phosphopeptide PSM counts at 1% FDR, site counts (excluding pA decoy sites) for all PSMs collapsed by peptidoform position and at each of the FLR thresholds; 1%, 5% and 10%.

| Dataset | 1% FDR<br>PSM<br>Count | 1% FDR<br>Phospho-<br>peptide<br>Count | Peptidofo<br>rm-site<br>count | 1% FLR<br>Peptidofo<br>rm Site<br>Count | 5% FLR<br>Peptidofo<br>rm Site<br>Count | 10% FLR<br>Peptidofo<br>rm Site<br>Count |
| --- | --- | --- | --- | --- | --- | --- |
| PXD002222 | 39092 | 29086 | 6762 | 1935 | 2951 | 3649 |
| PXD004939 | 81254 | 62935 | 10701 | 2156 | 4094 | 5348 |
| PXD005241 | 127696 | 106459 | 26190 | 1701 | 4687 | 8872 |
| PXD004705 | 80741 | 61029 | 10258 | 2050 | 3989 | 5133 |
| PXD002756 | 69232 | 18203 | 9734 | 1421 | 3708 | 5581 |
| PXD012764 | 45184 | 14020 | 5600 | 695 | 1269 | 1677 |
| PXD000923 | 14852 | 12167 | 5965 | 2366 | 3708 | 4221 |
| PXD019291 | 31094 | 30764 | 22128 | 4072 | 11034 | 13984 |

Different datasets contributed between ∼700 and ∼4,000 sites at the strictest FLR 1% threshold, and between 1,700 and ∼14,000 sites at 10% FLR. When performing PTM site localisation, there is a steep drop off in sensitivity when applying strict FLR thresholding (at say 1%), compared to weak thresholding (10% FLR) or say performing no explicit FLR thresholding – indicative of the challenge of confident site localization. Table 1 displays the counts of sites observed in the original studies and the statistical controls performed. Site counts ranged from ∼2,000 up to ∼9,000. In no original analysis (as published originally) was global FLR estimated, although this is not surprising since methods for accurate FLR estimation have not been well described until recently. While some *ad hoc* score thresholds were applied for local (i.e. per PSM) site scoring e.g. PhosphoRS > 0.9, this does not easily translate to a global (i.e. across the entire dataset) FLR, and thus it is reasonable to assume that there were variable (sometimes high) rates of FLR in different studies.

The tissues used for each dataset are shown in Table 1. These include flower, leaf, anther, shoot, panicles (young and mature), root and pollen samples. Although we can make no quantitative claims about site occupancy in a given tissue, by showing the tissues present in each dataset, a reader can infer if a given site has been seen in specific tissues.

We next performed a simple meta-analysis by combining all datasets, and assigned sites labels based on their scores and occurrences in datasets: Gold-Silver-Bronze (Table 3, Figure 1b). Decoy identifications of pAla were carried forward, enabling validation of the false reporting within these subsets. There are only two pAla hits within the Gold set, indicating that the overall FLR is very low in this subset. The meta-analysis also demonstrates that within each set, the reported counts for pTyr are relatively similar to pAla (taking into account that Ala is more abundant than Tyr in the proteome), indicating that pTyr hits reported for these datasets are likely to be mostly/entirely false positives, and should be treated with caution when interpreting any reported observations of pTyr from these datasets in rice. We recorded five unique Gold category pTyr sites. When looking at the scores of the spectra supporting these sites (Supplementary Figure 1), it was seen that most of these had only weak evidence supporting them and may be false positives.

**Table 3:** Count of sites (uniquely mapped to genomic loci), per category for Gold, Silver and Bronze. Totals target sites for categories are excluding alanine sites.

|  | **Gold** | **Silver** | **Bronze** | **Total** |
| --- | --- | --- | --- | --- |
| **A** | 2 | 53 | 420 | 475 |
| **S** | 2248 | 4397 | 7233 | 13878 |
| **T** | 212 | 499 | 850 | 1561 |
| **Y** | 9 | 25 | 92 | 126 |
| **Total target sites** | 2469 | 4921 | 8175 | 15565 |

In Figure 1c, we display the counts of sites depending in the source database – 13,425 sites were identified on peptides within proteins from all three databases (MSU, RAP-DB and UniProtKB). The original source of UniProtKB proteins is RAP-DB (with some later manual curation) – and we can observe 465 sites observed in RAP-DB and UniProtKB, but not in MSU, giving indications of peptides where the source RAP-DB gene model is likely superior to the MSU alternative. For the 212 sites that are common to MSU and UniProtKB, but not present in RAP-DB, would indicate that UniProtKB curators have altered gene models, such that they contain peptides identical to MSU. For 112 sites identified in MSU and RAP-DB, but not in UniProtKB, it is possible that UniProtKB curation has removed correct sections of gene models, or these sequences are entirely absent from UniProtKB. There are few phosphosites unique to RAP-DB or to UniProtKB, but 1,302 unique to MSU-derived protein sequences. The MSU annotation contains a larger count of protein sequences (48,237) than RAP-DB (46, 665), and many gene models different to the RAP-DB annotation. The identification of many phosphosites unique to MSU sequences, gives evidence for gene models that should be added or updated in the RAP-DB source.

In our mapping process from peptidoforms to proteins, we take the approach that if a peptide can be matched to proteins from multiple different locus, then all mappings should be accepted (unlike traditional proteomics approaches where parsimony in reporting protein identifications is preferred). The rationale is that the evidence presented is that a given peptidoform has been observed with a phosphosite, although due to the nature of tandem MS/MS, it is not possible to say definitely which protein was actually observed (when the peptidoforms matches multiple). If two proteins with highly similar sequences overall (and in this case an identical peptide sequence that has been identified), it seems probable that both can be phosphorylated on the identified position. The counts of phosphosites mapped onto one or more proteins is displayed in Figure 1d – the vast majority of sites are mapped to one or two proteins only, with a small count of sites mapped to multiple proteins, including 71 sites mapped to >=10 proteins. This happens in cases of very expanded gene families in rice, with paralogues of near identical sequence – it is not possible to determine which protein was actually observed in the experiment.

### Data visualisation

The gold-silver-bronze classified data has been loaded into UniProtKB for visualisation alongside other datasets and information available for rice. As one example, the phosphopeptides can be viewed in the context of AlphaFold2 (AF2) [56] predicted protein structures (Figure 2). The protein visualised in this case is OSCA1.2 (hyperosmolality-gated calcium-permeable channel 1.2, UniProtKB: Q5TKG1, MSU: LOC_Os05g51630, RAP-DB: Os05t0594700) and has a pSer at position 50. The AlphaFold prediction suggests that the serine forms a hydrogen bond with Arg36. It has been shown that phosphorylation can strengthen hydrogen bonds with Arg residues [57], and thus the pSer may have a structural role. With the widespread availability (now) of both phosphosite and structural data from AF2 models in UniProtKB, this provides a significant resource for rice cell signalling research.

**Figure 2:**
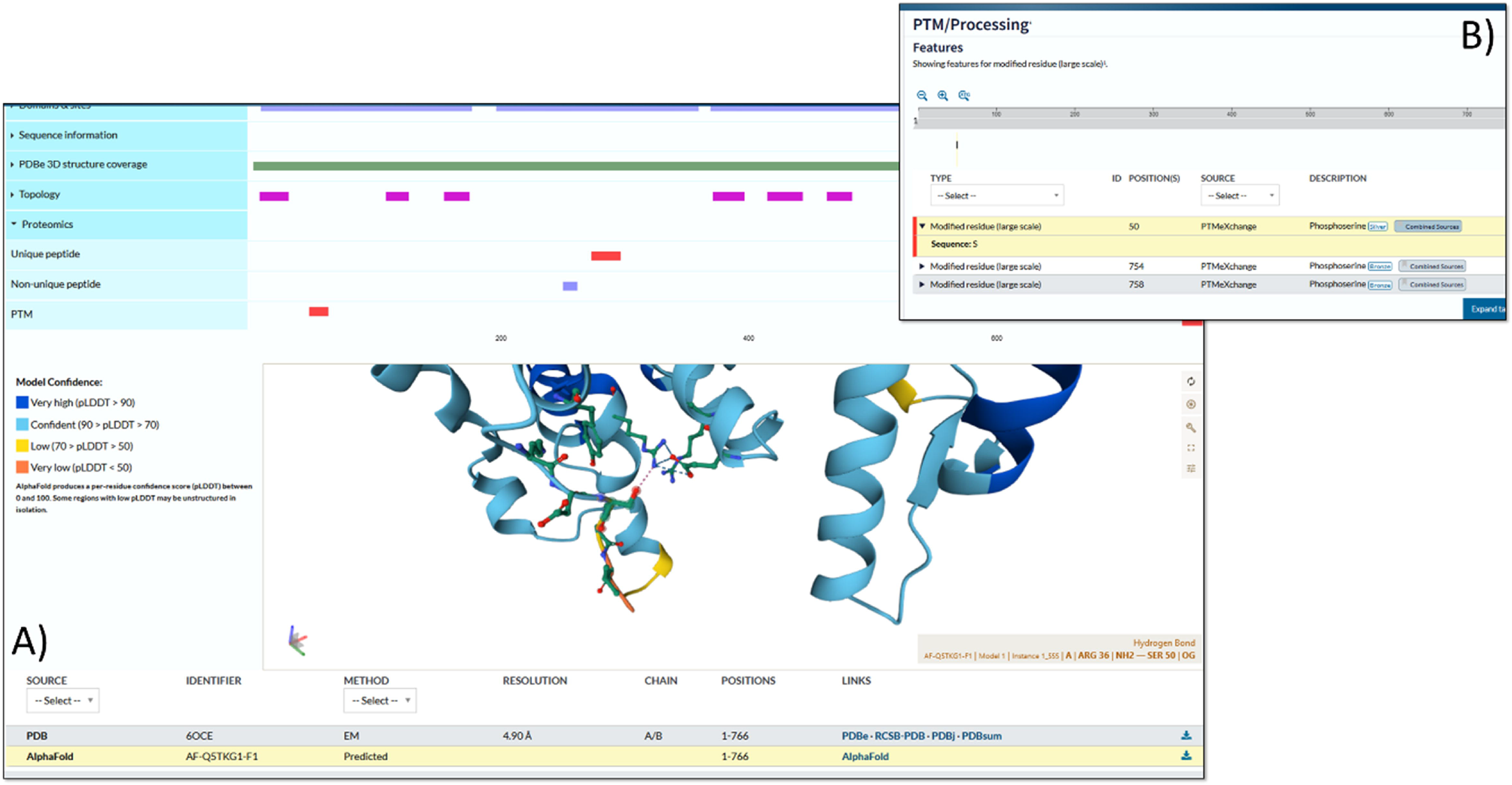
A display in UniProtKB of protein Q5TKG1 (RAP-DB: Os05t0594700-01; LOC_Os05g51630.1) showing A) three identified phosphosites in the tabular view and the B) structural context of the site on an AlphaFold2 prediction.

The build has also been loaded into PeptideAtlas, enabling browsing or searching the evidence for individual sites, peptides and proteins. An example of evidence supporting a phosphosite identified on a peptide is shown in the Supplementary materials (Supplementary Figure 2). We also demonstrate how Universal Spectrum Identifiers (USIs) can be used to visualise spectra supporting modification positions and can be a valuable tool to investigate the evidence supporting identified modifications (Supplementary Figure 3). The USI with the highest site probability for identified sites can be located in Supp Data File 3. Loading one of these USIs via https://proteomecentral.proteomexchange.org/usi/ imports the spectra and the claimed identification. By altering the position of the modification, it is possible to test which ions support an alternative hypothesis (site position in the peptide).

### Motif and pathway enrichment analysis

We ran motif analysis on the full set of identified pSer or pThr phosphosites (Gold+Silver+Bronze) using rmotifx (Figure 3), to identify enrichment of amino acids proximal to phosphosites potentially indicative of kinase families responsible for those sites (Supplementary Data File 1). Supplementary Figure 4 displays plots of the most enriched amino acids at each position, relative to the phosphosite for significant motifs. Numerous motifs are identified, with commonly enriched amino acids being P at +1 (relative to the target site), D/E at −1, +1, +2 and several others.

**Figure 3:**
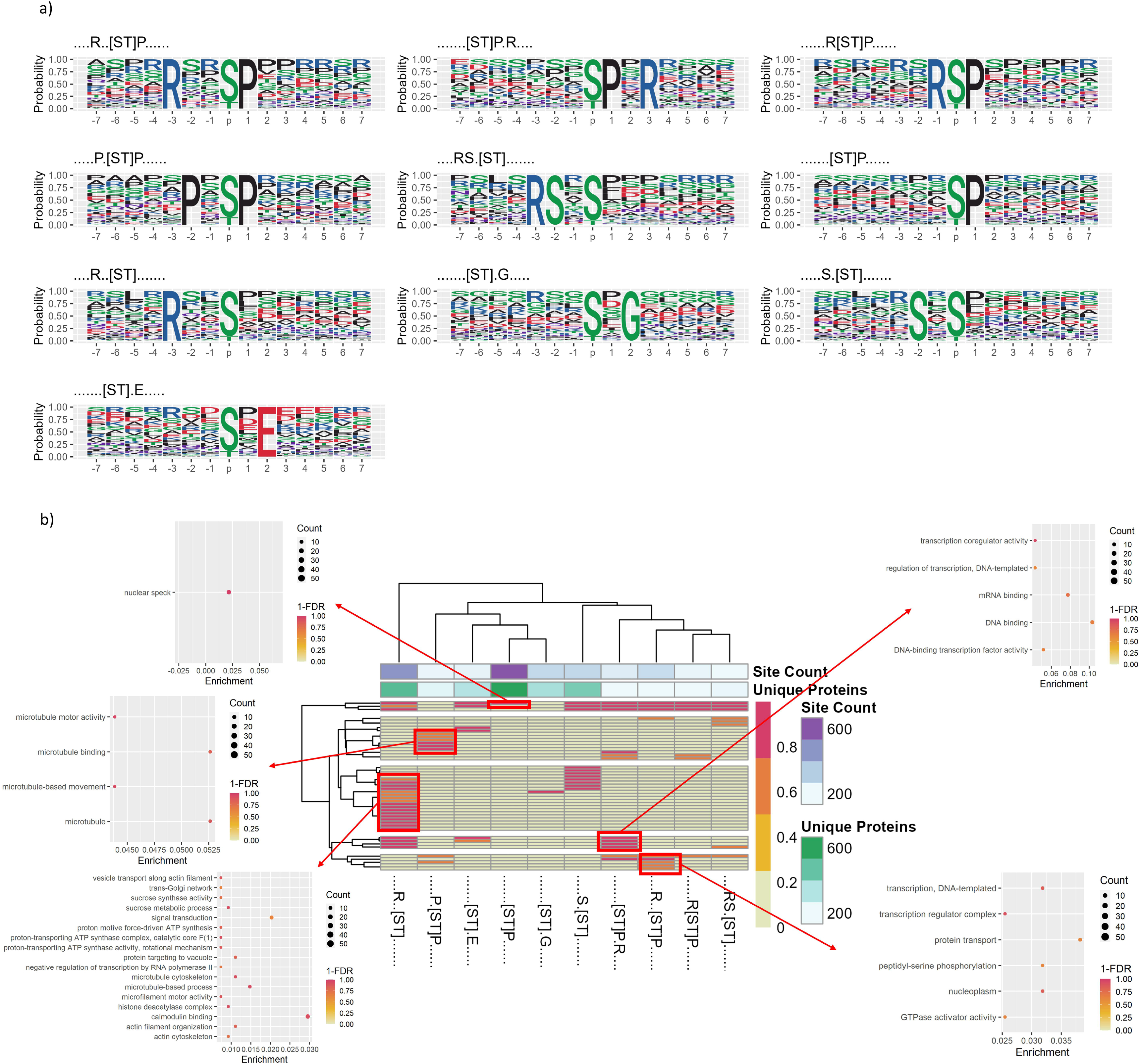
a) Motif logos showing the probability of particular amino acids to be present, surrounding the S/T phosphosites within Gold, Silver and Bronze datasets (motifs were filtered to be seen in at least 100 unique proteins). b) Heat map displaying significant motifs versus GO term clusters from Cluster Profiler (y-axis), displaying 1-FDR colour scale for pathway enrichment from proteins containing that phosphorylation, annotated with the count of unique proteins containing each motif and the count of phosphorylation sites supporting each motif.

There is a trade-off where having a higher overall count of true sites is likely to improve discovery of significant motifs, but too many false positive sites will weaken statistical power. As such, we also ran rmotifx analysis on “Gold+Silver” and “Gold only” sets (Supplementary Figures 4 and 5). Supplementary Table 2 displays a comparison of motifs discovered on different subsets of data (“Gold+Silver+Bronze” Gold+Silver” vs “Gold only”). The largest number of significant phosphorylation motifs was found for “Gold+Silver+Bronze”, which were thus used for the main analysis. We filtered those motifs to those found in at least 100 proteins, as shown in Figure 3a – indicating five motifs with proline (P) at the +1 position, four motifs with arginine (R) in a minus position (−1 or −3), and several others.

We next wished to explore whether such signatures related to differences in the pathways in which phosphorylated proteins act. We performed enrichment analysis to identify the pathways in which proteins containing significant motifs were acting (against a background of all rice phosphoproteins), using clusterProfiler (summarised in Figure 3b for motifs found in at least 100 proteins, results for all motifs shown in Supplementary Figure 6 and Supplementary Data File 2). Distinct enrichment of significant terms was obtained for different motifs. As examples, [ST]P.R motif-containing proteins were enriched for GO terms related to microtubules (“microtubule motor activity” (GO:0003777) and “microtubule-based movement” (GO:0007018)), compared to similar motif R..[ST]P was enriched for GO terms related to regulation of transcription (“transcription coregulator activity” (GO:0003712)), DNA and mRNA binding (“DNA polymerase III complex” (GO:0009360) and “mRNA splicing, via spliceosome” (GO:0000398)). P.[ST]P motif-containing proteins were enriched for “transcription regulator complex” (GO:0005667). R..[ST] motif-containing proteins were enriched in many GO terms, including “calmodulin binding” (GO:0005516), microtubule related terms, including “microtubule binding” (GO:0008017), “microtubule motor activity” (GO:0003777) and “microtubule cytoskeleton” (GO:0015630), amongst others), proton transport (“proton-transporting ATP synthase complex” (GO:0045261)) and “histone deacetylase complex” (GO:0000118). The rice kinome contains ∼1,500 kinases [58] much larger than the kinome in mammalian systems (humans have around 600 kinases for example). Even in humans, accurate assignment of kinase-substrate relationships is especially challenging, and for rice, given the sparsity of experimental data on kinase-substrate relationships, it is not possible to make accurate predictions about the kinases responsible for individual sites. However, the motif groups and downstream pathways identified here provided a starting point for interpreting the high-level different signalling pathways, which presumably correspond to different families of kinases.

### SAAV Analysis

We next assigned all phosphosites into five categories, determined in relationship to known non-synonymous SNPs (i.e. single amino acid variants – SAAVs) from the rice 3,000 genome set [59], whereby a category 1 site has an amino acid polymorphism in the reference genome (Nipponbare), causing a loss of this phosphosite in some other varieties (Figure 4). In the whole dataset, excluding pA decoy sites, there are 388 category 1 sites (Figure 4a), which are further explored in Figure 4b showing the most commonly substituted amino acid. Over-represented substitutions included S->L. Under-represented substitutions were D/E/H/K/Q/V. S/T->D mutations are potentially of great interest as Asp can mimic pSer/pThr as a constitutively active phosphorylation site, which could be a dominant allele for breeding. However, in our data, we saw only a single phosphosite (Bronze FLR category) with a S->D mutation (in LOC_Os10g32980.1 / “Cellulose synthase A7”). The implied amino acid substitution only observed at very low minor allele frequency (0.00033 i.e. one single cultivar in the 3,000 set), which could also be a sequencing error. We thus conclude that pSer-> Asp phospho-mimetic substitutions are exceedingly rare in the rice proteome. Within the data, there are 25 cases of S->T and 4 T->S phosphosite SAAVs (which would likely not disrupt phosphorylation).

**Figure 4.**
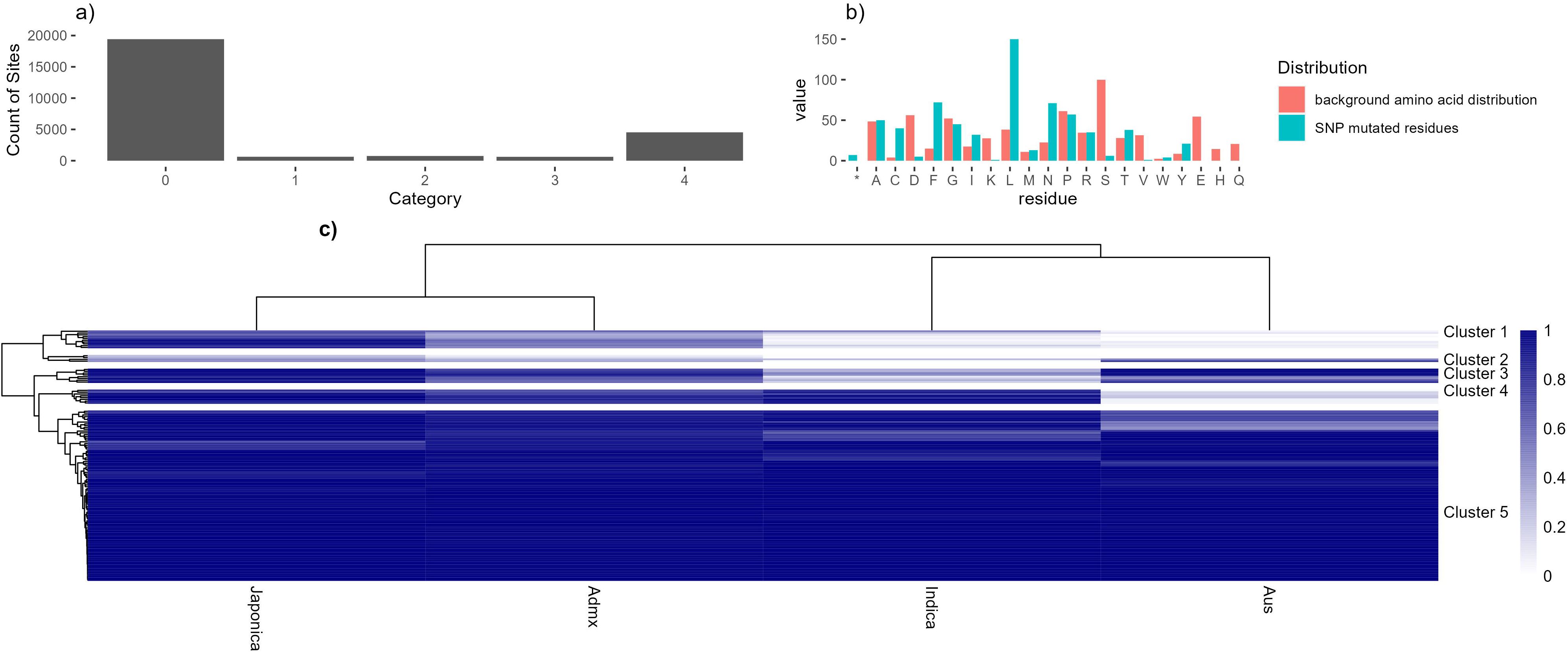
A) Bar chart showing counts of phosphosite by SAAV category; B) Counts of substituted amino acid in category 1 phosphosites, including the normalised background distribution of that amino acid (*=stop codon); C) A heat map to show the assumed major allele frequency (i.e. frequency of Ser / Thr within the 3K “pseudo-proteins” from the rice 3K SNP set) of the phosphosite in four rice family groups: Japonica, admixed, Indica and Aus. Allele frequencies are filtered for total difference between families >0.05 to remove genes showing the same frequency across all families.

**Figure 5.**
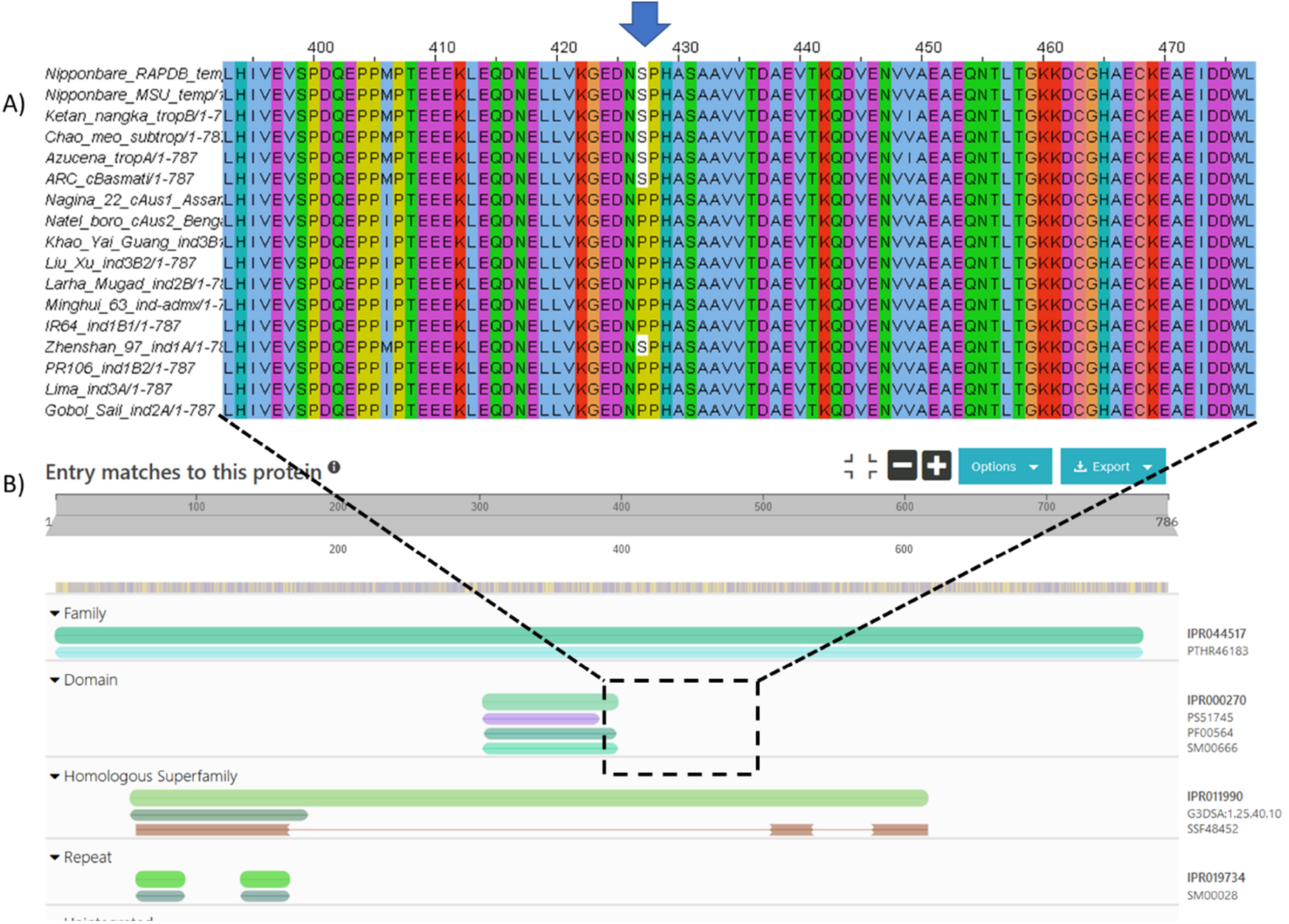
A) Protein sequence alignment of Os09g0135400 (RAP-DB), LOC_Os09g04990 (MSU) alongside protein sequences from other recently Oryza sativa varieties. B) The results of searching the protein sequence in InterProScan. The pSer site is nearby to PB1 protein binding domain (IPR000270, InterPro), and the protein is part of the Tetratricopeptide-like helical domain superfamily (IPR011990).

We also note five observations where the pSer has apparently been substituted with a stop codon (*) in proteins: LOC_Os03g17084.1 (Gold site on position 21, RAP-DB ID=Os03t0279000, annotated as “Similar to Histone H2B.1”; LOC_Os02g52780.1 (Gold site on position 98, RAP-DB ID=Os02t0766700-01, annotated as “BZIP transcription factor”), Bronze sites are seen on proteins LOC_Os02g07420.1, LOC_Os05g39730.1 and LOC_Os01g31010.1. However, in all cases, SNPs were at very low minor allele frequencies (1 – 3 cultivars only out of the entire 3,000 set), indicating that phosphosite mutation to a stop codon is an exceptionally rare event in the rice pan genome. Supplementary Data File 3 shows the genomic location of all identified phosphorylation sites along with the SAAV positions and gene annotations.

We converted category 1 SAAV data into a heat map (Figure 4C), with clustering of sites by major allele frequency in four rice families: Japonica, Indica, Admixed and Aus, with a tree cut method to split the dendrogram into sub-groups (Supplementary Data File 4). Five distinct clusters can be observed (as labelled). 1) high in Japonica and admixed, low in Indica and Aus families; 2) variable pattern cluster, mostly medium to low conservation in all families; 3) high in three families, lower in Indica; 4) high in three families, low in Aus; and 5) high in all families. Supplementary Data File 4 contains the source data, enabling the data to be filtered to find alleles of interest, where there are likely significant differences between major varietal groups. Cluster 4 (Table 4) contains phosphosites that are mostly conserved in Japonica, Indica and admixed varieties, but lowly conserved in Aus. Aus variety rice cultivars are generally considered to be resistant to biotic stress, like drought. Source genes mostly have limited annotation, although genes with annotations include part of the Tho complex (involved with mRNA transcription, processing and nuclear export) and a Zinc finger protein (members within this large family of proteins have been implicated in transcriptional regulation and responses to stress). Phosphosites in proteins lacking annotation may be good candidates for further study, for potential roles in Aus-specific phenotypic responses.

**Table 4.**
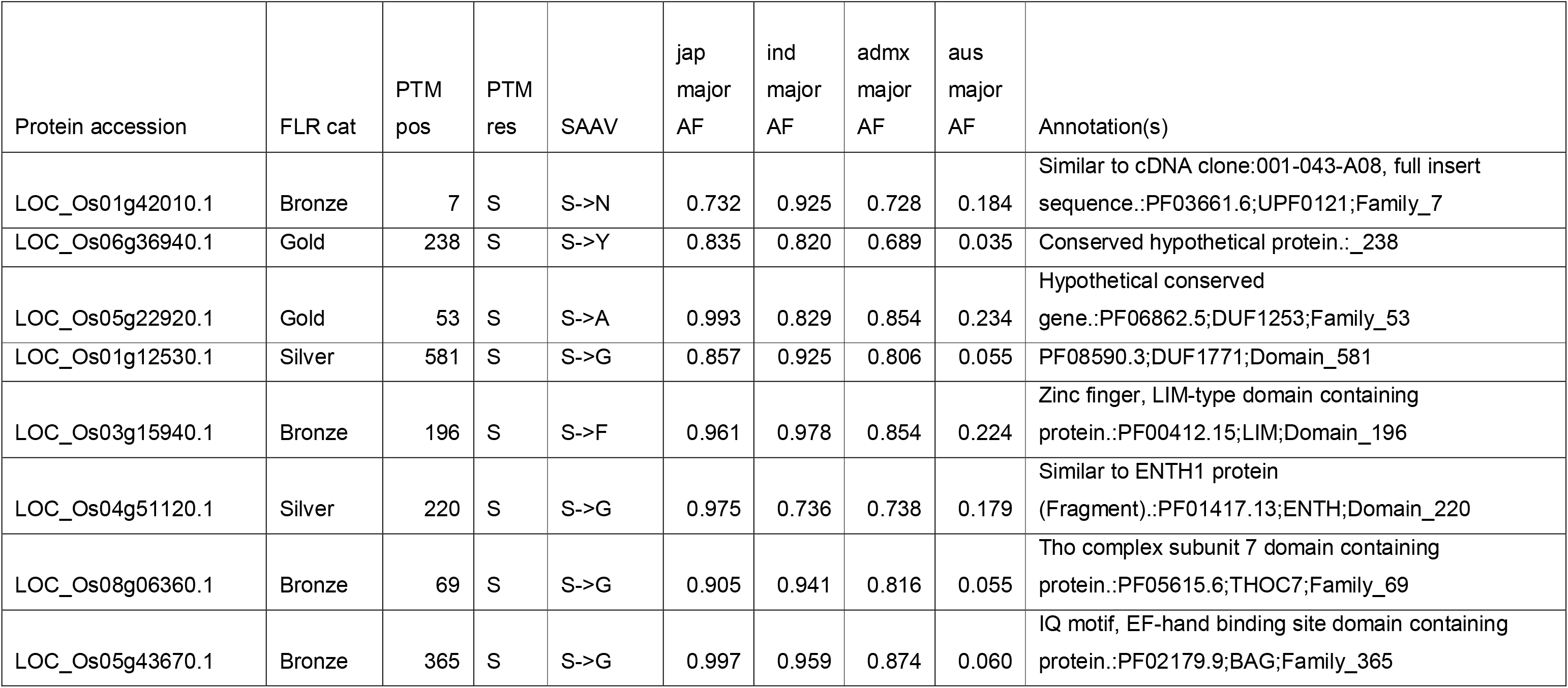
Phosphosites identified in cluster 4 on Figure 4, defined by the pattern of major allele frequencies (AF) across four varietal groups – Japonica (jap), Indica (ind), admixed (admx) and aus. Annotations are sourced by merging any data held in MSU or RAP-DB databases for the corresponding gene. Cluster 4 is mostly characterised by high AF in Japonica, Indica, admixed, but low in Aus.

Cluster 1 phosphosites are those present at high allele frequencies in Japonica and admixed but lower in Indica and Aus type rices – indicating potential cell signalling differences across the two major branches of *Oryza sativa* (summarised in Table 5). Proteins of potential interest for further study include LGD1 (“Lagging Growth and Development”), “Cullin 1” (LOC_Os03g44900.1), and HSP40. LGD1 has been implicated in regulation of plant growth and yield [60].

**Table 5.**
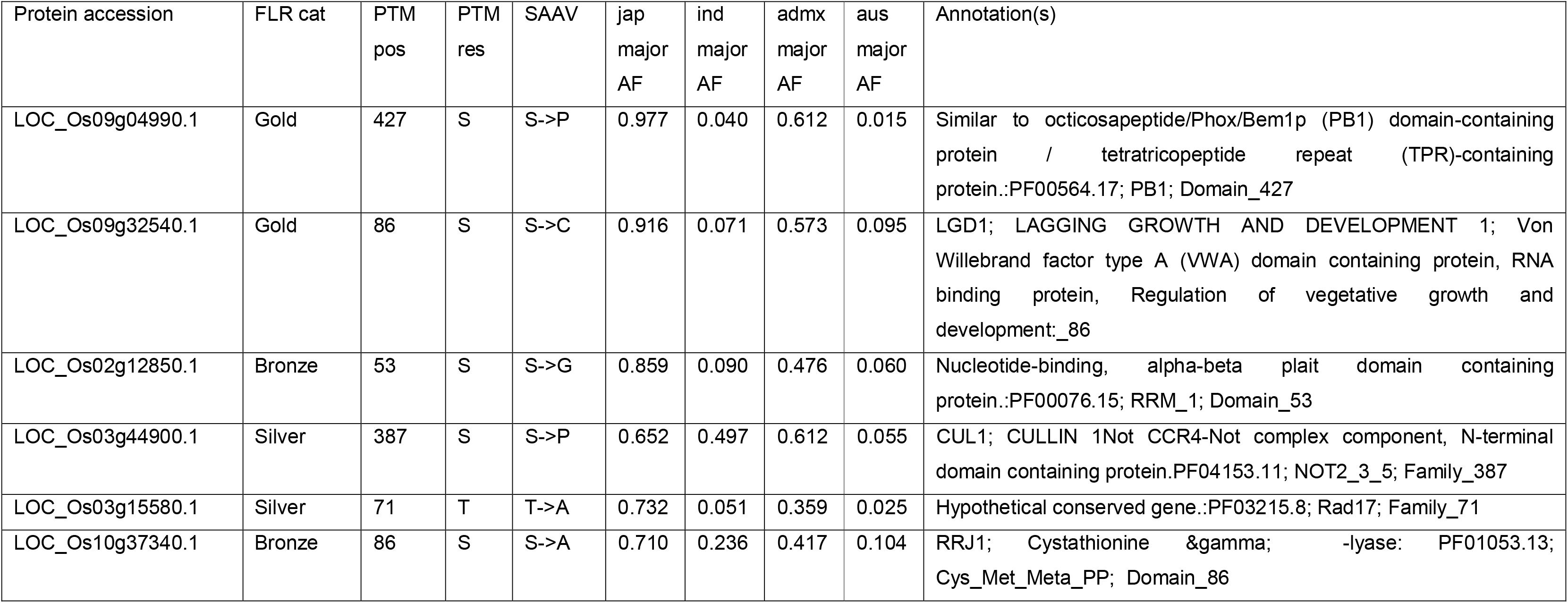
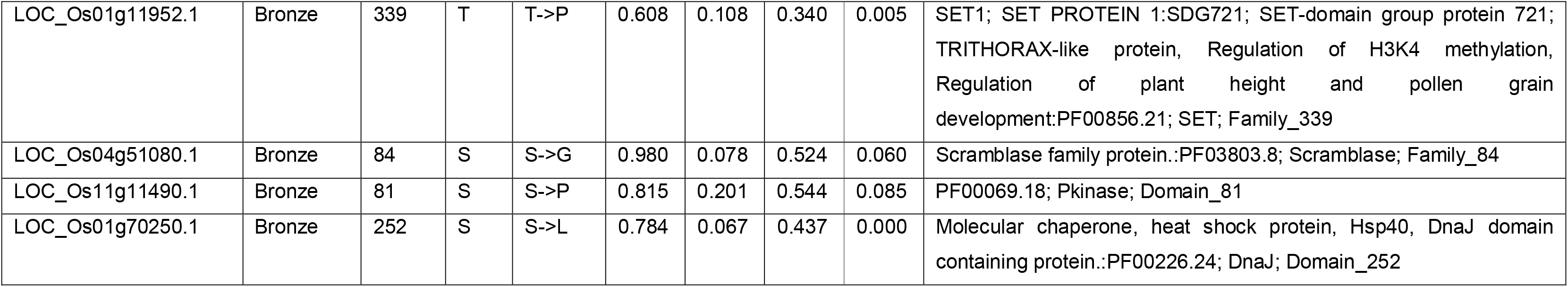
Phosphosites identified in cluster 1 on Figure 4, defined by the pattern of major allele frequencies (AF) across four varietal groups – Japonica (jap), Indica (ind), admixed (admx) and aus. Annotations are sourced by merging any data held in MSU or RAP-DB databases for the corresponding gene. Cluster 1 is mostly characterised by high AF in Japonica, and admixed, but low in Aus and Indica.

The gene annotated in OryzaBase as OsCullin1 (LOC_Os03g44900.1) appears to have been misnamed in this publication [61], based on apparent shared homology to *Arabidopsis thaliana* Cullin 1 (annotated in TAIR [62] to act as a component of ubiquitin ligase, with roles in response to auxin and jasmonic acid). However, LOC_Os03g44900.1 has high homology to NOT family transcriptional regulators, and has characteristic domains of this family, and should be renamed in OryzaBase.

The SAAV analysis presented above generated “pseudo-protein” sequences by substituting amino acids, based on short DNA read data from the 3K rice genome set, which have been mapped against the Nipponbare reference genome. There is thus potential for assumed SAAVs to be incorrect, due to sequencing errors (as the 3K set does not always have high depth of coverage), if gene structure genuinely differs across different varietal groups, or if the RAP-DB or MSU gene model for Nipponbare is not correct. To validate the SAAV data, we also mapped the phosphosites to recently released gene models for 16 new rice varieties, called the “MAGIC-16” [63]. In Figure 6, we display the protein sequence alignment across orthologs for cluster 1 protein LOC_Os09g04990.1 (Os09g0135400) position 427, with the position of an identified phosphosite marked. It can be observed that the pSer is present in tropical, sub-tropical, temperate and aromatic varieties, but absent in all indica varieties, except Zhenshan 97. The equivalent allele frequencies for this pSer site are trop_ref_freq=0.99; temp_ref_freq=0.99; admix_ref_freq=0.61; japx_ref_freq=0.99; subtrop_ref_freq=0.97; aus_ref_freq=0.01; aro_ref_freq=0.93; ind2_ref_freq=0.02; indx_ref_freq=0.05; ind1B_ref_freq=0.08; ind3_ref_freq=0.01; ind1A_ref_freq=0.03 – which appears to be in-line with the genuine protein sequences from the MAGIC-16 set. Protein sequences for all the MAGIC-16 set are available from Ensembl Plants and Gramene, enabling any phosphosites identified in this resource, to be cross-referenced to protein sequences annotated from high-quality whole genome assemblies, prior to any experimental work being conducted to validate PTM site differences.

## Conclusions

In this work, we have performed a meta-analysis of phosphoproteomics datasets for rice, mapped onto the reference Nipponbare proteome. The pipeline includes conservative statistics to avoid reporting false positives, and a simple Gold-Silver-Bronze metric allowing users of the data to focus understand the likelihood of a site being correct.

The dataset has been deposited into UniProtKB enabling sites to be analysed alongside any other data held there about protein structure/function, including AlphaFold2 predictions for all rice proteins. The data is also available in PeptideAtlas and PRIDE, enabling detailed exploration of scores and visualization of source mass spectra, as a full evidence trail.

We have also mapped the data to variation coming from the 3,000 genome set, creating a resource for allele mining, where phosphosites are likely to have lost function due to amino acid substitutions in some rice varieties, with alterations to downstream cell signalling pathways. We expect this will be a powerful resource for rice biology, and all datasets are fully open and available for re-analysis.

## Supplemental Data

**Supplementary Data File 1 – All_motifs**: motifs found in each of the three categories (“Gold, Silver and Bronze”, “Gold and Silver” and “Gold only”) with enrichment scores and protein counts.

**Supplementary Data File 2 - Genomic_site_data_w_SNP_annotation**: All phosphosites identified in the study, along with their mapped genomic position, data on SNPs, single amino acid variants and functional annotations.

**Supplementary Data File 3 - ClusterProfiler GO enrichment GSB motifs**: Enriched GO terms for each motif identified around phosphosites scored in the “Gold, Silver and Bronze” category.

**Supplementary Data File 4 - Figure_4b_heatmap_clusters**: Clusters of genes seen in the heatmap shown in figure 4b.

## Acknowledgements

We are grateful to all the groups that generated MS data and deposited into PRIDE/ProteomeXchange to allow for re-analyses, and those who generated the 3K and MAGIC-16 rice resources.

## Author Contributions

K.A.R. performed MS data curation, searching, analysis and manuscript writing. A.P. supported MS data curation. Y.P.R. supported PRIDE data upload and curation. O.M.C. facilitated statistical data analysis. D.J.K. assisted with metadata dataset curation. E.B-B, M.M. and J.F. assisted with data loading into UniProtKB. D.C. and K.L.M. assisted with rice 3K data analysis. E.W.D assisted with MS data searching and project supervision. Z.S. assisted with MS data search and the loading into PeptideAtlas. J.A.V. assisted with PRIDE data loading and project supervision. A.R.J. coordinated the research, assisted with data analysis and writing the manuscript.

## References

1. Fornasiero, A., R.A. Wing, and P. Ronald, Rice domestication. Curr Biol, 2022. 32(1): p. R20–R24.

2. International Rice Genome Sequencing, P., The map-based sequence of the rice genome. Nature, 2005. 436(7052): p. 793–800.

3. Yu, J., et al., A draft sequence of the rice genome (Oryza sativa L. ssp. indica). Science, 2002. 296(5565): p. 79–92.

4. Koizumi, T., S.H. Gay, and G. Furuhashi, Reviewing Indica and Japonica rice market developments. OECD Food, Agriculture and Fisheries Papers, No. 154, 2021.

5. Ouyang, S., et al., The TIGR Rice Genome Annotation Resource: improvements and new features. Nucleic Acids Res, 2007. 35(Database issue): p. D883–7.

6. Sakai, H., et al., Rice Annotation Project Database (RAP-DB): an integrative and interactive database for rice genomics. Plant Cell Physiol, 2013. 54(2): p. e6.

7. Tello-Ruiz, M.K., P. Jaiswal, and D. Ware, Gramene: A Resource for Comparative Analysis of Plants Genomes and Pathways. Methods Mol Biol, 2022. 2443: p. 101–131.

8. Yates, A.D., et al., Ensembl Genomes 2022: an expanding genome resource for non-vertebrates. Nucleic Acids Res, 2022. 50(D1): p. D996–D1003.

9. UniProt, C., UniProt: the Universal Protein Knowledgebase in 2023. Nucleic Acids Res, 2023. 51(D1): p. D523–D531.

10. Wang, W., et al., Genomic variation in 3,010 diverse accessions of Asian cultivated rice. Nature, 2018. 557(7703): p. 43–49.

11. Zhou, Y., et al., A platinum standard pan-genome resource that represents the population structure of Asian rice. Sci Data, 2020. 7(1): p. 113.

12. Song, J.M., et al., Two gap-free reference genomes and a global view of the centromere architecture in rice. Mol Plant, 2021. 14(10): p. 1757–1767.

13. Blanco-Tourinan, N., A. Serrano-Mislata, and D. Alabadi, Regulation of DELLA Proteins by Post-translational Modifications. Plant Cell Physiol, 2020. 61(11): p. 1891–1901.

14. Orosa-Puente, B., et al., Root branching toward water involves posttranslational modification of transcription factor ARF7. Science, 2018. 362(6421): p. 1407–1410.

15. Rigbolt, K.T. and B. Blagoev, Quantitative phosphoproteomics to characterize signaling networks. Semin Cell Dev Biol, 2012. 23(8): p. 863–71.

16. Ramsbottom, K.A., et al., Method for Independent Estimation of the False Localization Rate for Phosphoproteomics. J Proteome Res, 2022. 21(7): p. 1603–1615.

17. Camacho, O.M., et al., Assessing Multiple Evidence Streams to Decide on Confidence for Identification of Post-Translational Modifications, within and Across Data Sets. J Proteome Res, 2023. 22(6): p. 1828–1842.

18. Kalyuzhnyy, A., et al., Profiling the Human Phosphoproteome to Estimate the True Extent of Protein Phosphorylation. J Proteome Res, 2022. 21(6): p. 1510–1524.

19. Willems, P., et al., The Plant PTM Viewer, a central resource for exploring plant protein modifications. Plant J, 2019. 99(4): p. 752–762.

20. Heazlewood, J.L., et al., PhosPhAt: a database of phosphorylation sites in Arabidopsis thaliana and a plant-specific phosphorylation site predictor. Nucleic Acids Res, 2008. 36(Database issue): p. D1015–21.

21. Perez-Riverol, Y., et al., The PRIDE database resources in 2022: a hub for mass spectrometry-based proteomics evidences. Nucleic Acids Res, 2022. 50(D1): p. D543–D552.

22. Deutsch, E.W., et al., The ProteomeXchange consortium in 2020: enabling ‘big data’ approaches in proteomics. Nucleic Acids Res, 2020. 48(D1): p. D1145–D1152.

23. Desiere, F., et al., The PeptideAtlas project. Nucleic Acids Res, 2006. 34(Database issue): p. D655–8.

24. van Wijk, K.J., et al., The Arabidopsis PeptideAtlas: Harnessing worldwide proteomics data to create a comprehensive community proteomics resource. Plant Cell, 2021. 33(11): p. 3421–3453.

25. Vizcaino, J.A., et al., ProteomeXchange provides globally coordinated proteomics data submission and dissemination. Nat Biotechnol, 2014. 32(3): p. 223–6.

26. Martens, L., et al., PRIDE: the proteomics identifications database. Proteomics, 2005. 5(13): p. 3537–45.

27. Wang, K., et al., Analysis of phosphoproteome in rice pistil. Proteomics, 2014. 14(20): p. 2319–34.

28. Lu, Q., et al., Identification and characterization of chloroplast casein kinase II from Oryza sativa (rice). J Exp Bot, 2015. 66(1): p. 175–87.

29. Hou, Y., et al., A comprehensive quantitative phosphoproteome analysis of rice in response to bacterial blight. BMC Plant Biol, 2015. 15: p. 163.

30. Li, M., et al., Proteomic Analysis of Phosphoproteins in the Rice Nucleus During the Early Stage of Seed Germination. J Proteome Res, 2015. 14(7): p. 2884–96.

31. Qiu, J., et al., A Comprehensive Proteomic Survey of ABA-Induced Protein Phosphorylation in Rice (Oryza sativa L.). Int J Mol Sci, 2017. 18(1).

32. Wang, Y., et al., A phosphoproteomic landscape of rice (Oryza sativa) tissues. Physiol Plant, 2017. 160(4): p. 458–475.

33. Hou, Y., et al., A Quantitative Proteomic Analysis of Brassinosteroid-induced Protein Phosphorylation in Rice (Oryza sativa L.). Front Plant Sci, 2017. 8: p. 514.

34. Ye, J., et al., Proteomic and phosphoproteomic analyses reveal extensive phosphorylation of regulatory proteins in developing rice anthers. Plant J, 2015. 84(3): p. 527–44.

35. Yang, J., et al., Phosphoproteomic Profiling Reveals the Importance of CK2, MAPKs and CDPKs in Response to Phosphate Starvation in Rice. Plant Cell Physiol, 2019. 60(12): p. 2785–2796.

36. Al-Momani, S., et al., Comparative qualitative phosphoproteomics analysis identifies shared phosphorylation motifs and associated biological processes in evolutionary divergent plants. J Proteomics, 2018. 181: p. 152–159.

37. Inomata, T., et al., Proteomics Analysis Reveals Non-Controlled Activation of Photosynthesis and Protein Synthesis in a Rice npp1 Mutant under High Temperature and Elevated CO(2) Conditions. Int J Mol Sci, 2018. 19(9).

38. He, Z., et al., An L-type lectin receptor-like kinase promotes starch accumulation during rice pollen maturation. Development, 2021. 148(6).

39. Dai, C., et al., A proteomics sample metadata representation for multiomics integration and big data analysis. Nat Commun, 2021. 12(1): p. 5854.

40. Moosa, J.M., et al., Repeat-Preserving Decoy Database for False Discovery Rate Estimation in Peptide Identification. J Proteome Res, 2020. 19(3): p. 1029–1036.

41. Deutsch, E.W., et al., Trans-Proteomic Pipeline, a standardized data processing pipeline for large-scale reproducible proteomics informatics. Proteomics Clin Appl, 2015. 9(7-8): p. 745–54.

42. Deutsch, E.W., et al., Trans-Proteomic Pipeline: Robust Mass Spectrometry-Based Proteomics Data Analysis Suite. J Proteome Res, 2023. 22(2): p. 615–624.

43. Eng, J.K., T.A. Jahan, and M.R. Hoopmann, Comet: an open-source MS/MS sequence database search tool. Proteomics, 2013. 13(1): p. 22–4.

44. Keller, A., et al., Empirical statistical model to estimate the accuracy of peptide identifications made by MS/MS and database search. Anal Chem, 2002. 74(20): p. 5383–92.

45. Shteynberg, D., et al., iProphet: multi-level integrative analysis of shotgun proteomic data improves peptide and protein identification rates and error estimates. Mol Cell Proteomics, 2011. 10(12): p. M111 007690.

46. Shteynberg, D.D., et al., PTMProphet: Fast and Accurate Mass Modification Localization for the Trans-Proteomic Pipeline. J Proteome Res, 2019. 18(12): p. 4262–4272.

47. Jones, A.R., et al., The mzIdentML data standard for mass spectrometry-based proteomics results. Mol Cell Proteomics, 2012. 11(7): p. M111 014381.

48. Deutsch, E.W., et al., Universal Spectrum Identifier for mass spectra. Nat Methods, 2021. 18(7): p. 768–770.

49. R Core Team, R: A language and environment for statistical computing. 2021, R Foundation for Statistical Computing: Vienna, Austria.

50. Wagih, O., et al., Uncovering Phosphorylation-Based Specificities through Functional Interaction Networks. Mol Cell Proteomics, 2016. 15(1): p. 236–45.

51. Wu, T., et al., clusterProfiler 4.0: A universal enrichment tool for interpreting omics data. Innovation (Camb), 2021. 2(3): p. 100141.

52. Kurata, N. and Y. Yamazaki, Oryzabase. An integrated biological and genome information database for rice. Plant Physiol, 2006. 140(1): p. 12–7.

53. Zhang, J., et al., Extensive sequence divergence between the reference genomes of two elite indica rice varieties Zhenshan 97 and Minghui 63. Proc Natl Acad Sci U S A, 2016. 113(35): p. E5163–71.

54. Stein, J.C., et al., Genomes of 13 domesticated and wild rice relatives highlight genetic conservation, turnover and innovation across the genus Oryza. Nat Genet, 2018. 50(2): p. 285–296.

55. Contreras-Moreira, B., et al., GET_PANGENES: calling pangenes from plant genome alignments confirms presence-absence variation. Genome Biol, 2023. 24(1): p. 223.

56. Jumper, J., et al., Highly accurate protein structure prediction with AlphaFold. Nature, 2021. 596(7873): p. 583–589.

57. Mandell, D.J., et al., Strengths of hydrogen bonds involving phosphorylated amino acid side chains. J Am Chem Soc, 2007. 129(4): p. 820–7.

58. Dardick, C., et al., The rice kinase database. A phylogenomic database for the rice kinome. Plant Physiol, 2007. 143(2): p. 579–86.

59. Alexandrov, N., et al., SNP-Seek database of SNPs derived from 3000 rice genomes. Nucleic Acids Res, 2015. 43(Database issue): p. D1023–7.

60. Thangasamy, S., et al., Rice LGD1 containing RNA binding activity affects growth and development through alternative promoters. Plant J, 2012. 71(2): p. 288–302.

61. Zhao, J., et al., DWARF3 participates in an SCF complex and associates with DWARF14 to suppress rice shoot branching. Plant Cell Physiol, 2014. 55(6): p. 1096–109.

62. Berardini, T.Z., et al., The Arabidopsis information resource: Making and mining the “gold standard” annotated reference plant genome. Genesis, 2015. 53(8): p. 474–85.

63. Zhou, Y., et al., Pan-genome inversion index reveals evolutionary insights into the subpopulation structure of Asian rice. Nat Commun, 2023. 14(1): p. 1567.

