## Supplementary Figures and Tables for "A meta-analysis of rice phosphoproteomics data to understand variation in cell signalling across the rice pan-genome"

Supplementary Table 1: Supplementary Table 1: Summary of all rice phosphoproteomics datasets identified via the PRIDE repository.

| PXD | Title | PMID | Instrument | Count of .RAW files | Included in analysis? |
| --- | --- | --- | --- | --- | --- |
| PXD000857 | 29RNP phosphorylation site - Identification and characterization of chloroplast casein kinase II from Oryza sativa (rice) | 25316064 | LTQ Orbitrap | 1 | N |
| PXD000923 | Rice Pistil LC-MSMS - Analysis of Phosphoproteome in Rice Pistil | 25074045 | TripleTOF 5600 | 10 | Y |
| PXD001168 | 29RNP kinase - Identification and characterization of chloroplast casein kinase II from Oryza sativa (rice) | 25316064 | LTQ Orbitrap | 6 | N |
| PXD001774 | Un-conditioned commercial embryo culture media contain a large variety of non-declared proteins: a comprehensive proteomics analysis | 25164020 | TripleTOF 5600 |  | N |
| PXD002222 | Rice-xoo interaction phosphoproteome LC-MSMS | 26112675 | Q Exactive | 6 | Y |
| PXD002756 | Proteomic and Phosphoproteomic Analyses Reveal Extensive Phosphorylation of Regulatory Proteins in Developing Rice Anthers | 26360816 | LTQ Orbitrap Elite | 5 | Y |
| PXD004705 | A phosphoproteomic analysis of ABA induced protein phosphorylation revealed that Serine 71 is a major phosphosite for bZIP72 in SAPK10-mediated ABA signaling | 28439285 | Q Exactive | 9 | Y |
| PXD004939 | A comprehensive proteomic survey of ABA-induced protein phosphorylation in rice (Oryza sativa L.) | 28054942 | Q Exactive | 9 | Y |
| PXD005241 | A phosphoproteomic landscape of rice (Oryza sativa L.) tissues | 28382632 | Q Exactive | 18 | Y |
| PXD007979 | Comparative Qualitative Phosphoproteomics Analysis Identifies Shared Phosphorylation Motifs and Associated Biological Processes in Flowering Plants | 29654922 | TripleTOF 5600  LTQ Orbitrap Velos  Q Exactive | 124 | N |
| PXD010565 | Analysis of chloroplast phosphoproteome | Pre-publication | LTQ Orbitrap | 24 | N |
| PXD012764 | Phosphoproteomic profiling responds to phosphate starvation in rice | 31424513 | Q Exactive | 6 | Y |
| PXD019291 | ABNORMAL POLLEN 1 (AP1) encodes a legume-type lectin receptor-like kinase and is involved in starch accumulation during rice pollen maturation | 33658224 | Q Exactive | 4 | Y |

**Peptide Atlas dataset deposition and visualisation**

Localisation probabilities can be viewed through PeptideAtlas (<https://peptideatlas.org/builds/rice/phospho/>) in the form of histograms. Supplementary Figure 1a shows the histogram of the peptide “IAVQNAETLSLASPR”. Here we can see the distribution of scores around each potential phosphorylation site within the peptide. The serine at position 13 shows the best score distribution with 63 PSMs supporting this site within the scores 0.95<p<0.99 and 128 PSMs with scores 0.81<p<0.95. These supporting sites are summarised in Supplementary Figure 1b, with an example spectra for this peptide being displayed in Supplementary Figure 2a for the PSM with the Universal Spectrum Identifier “mzspec:PXD004939:Rice_phos_Nip_0h_20per_F1_R3:scan:23826:IAVQNAETLSLAS[Phospho]PR/2”. Supplementary Figure 2B shows how the annotation of the alternative position spectrum changes, highlighting how valuable USIs can be when investigating modification positions.

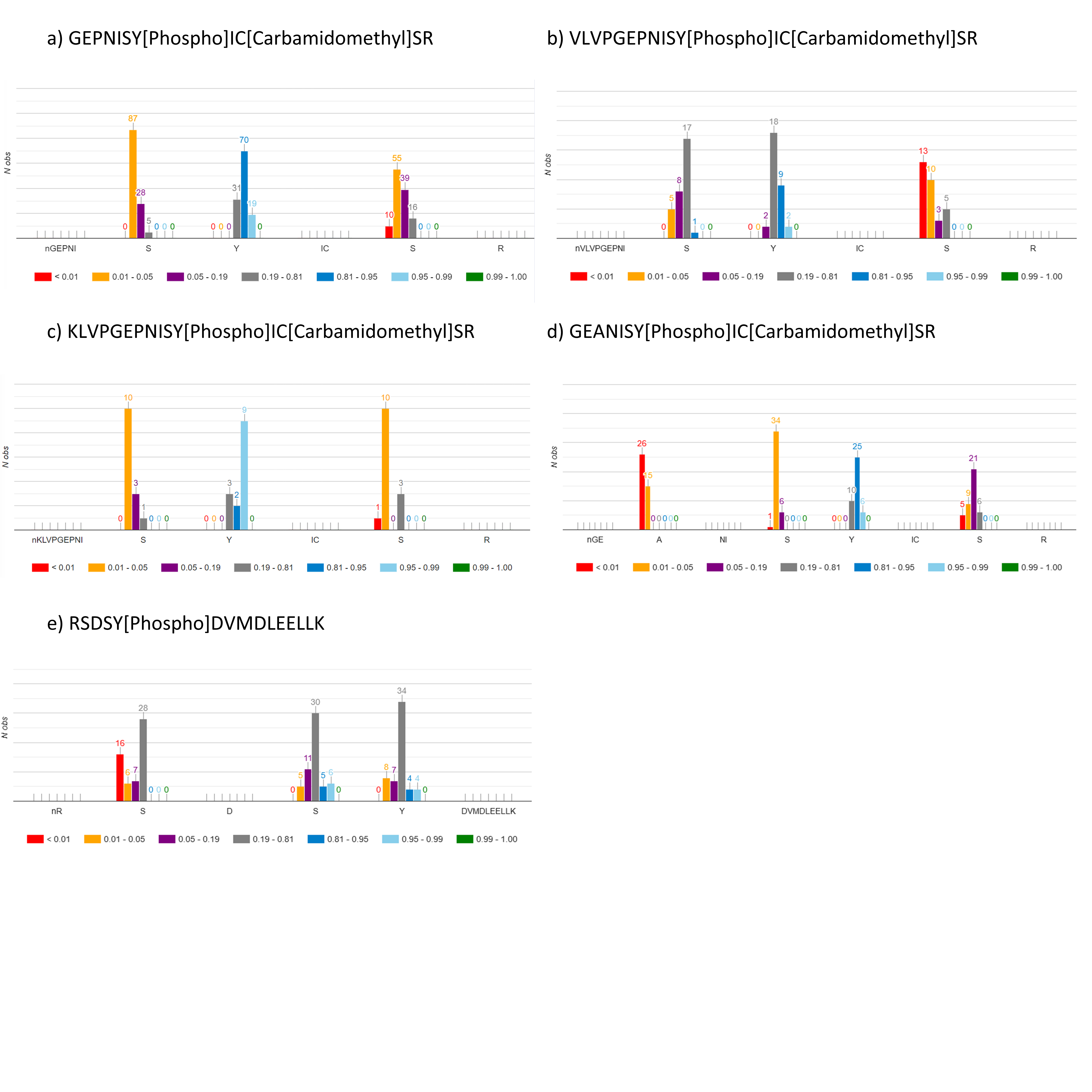

Supplementary Figure 1: Histograms showing scores of potential phosphosites on five peptidoforms identified with pTyr Gold scored sites. a) shows some evidence of phosphorylation of the Tyr site with 70 spectra falling within the “0.81-0.95” and 19 spectra in the “0.95-0.99” high confidence categories, showing that this could potentially be a true identification , b) shows some weak evidence for the Tyr site with 9 spectra in the “0.81-0.95” and 2 spectra in the “0.95-0.99” categories, c) shows some weak evidence for the Tyr site with 9 spectra being seen within the “0.95-0.99” score bin, d ) shows some fairly strong evidence of the Tyr site with 25 spectra in the “0.81-0.95” and 6 within the “0.95-0.99” high confidence score categories and e) shows weak evidence for the Y with 4 spectra within the “0.81-0.95” and 4 spectra in the “0.95-0.99” score ranges, with similar evidence being seen for the Ser at position 4, with 5 spectra within the “0.81-0.95” and 6 spectra in the “0.95-0.99” score ranges . Refer to Supplementary Figure 2 for more detail on Peptide Atlas build.

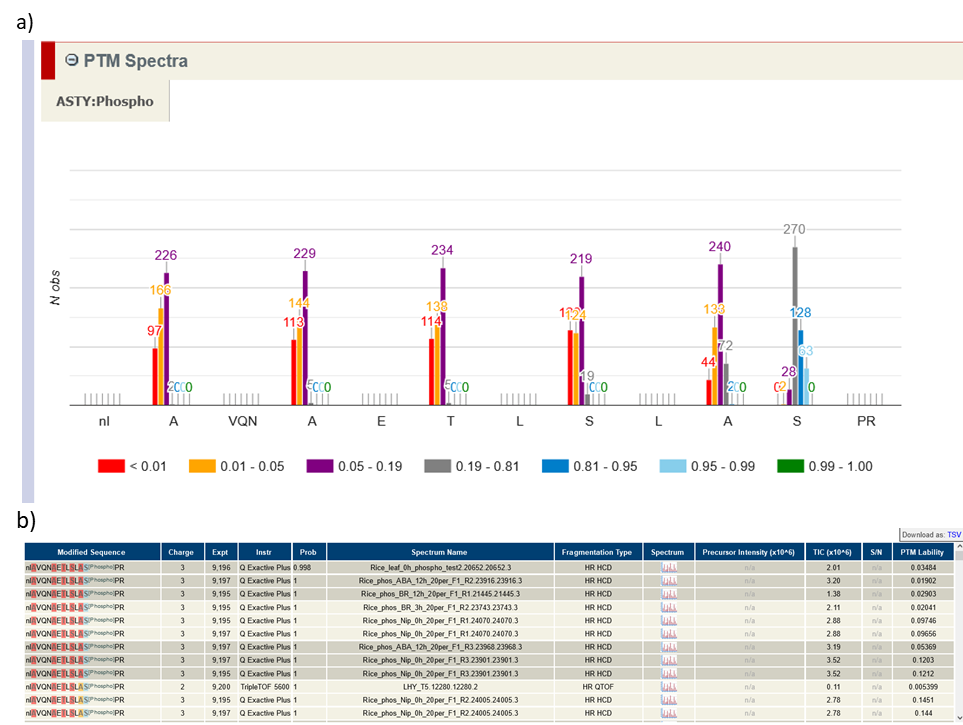

Supplementary Figure 2: a) Score histograms of peptide modification positions for peptide “IAVQNAETLSLASPR”, showing favouring of the serine at position 16 with more confident score distributions seen at this position than other potential sites. b) Summary table showing spectra supporting the peptide shown in a).

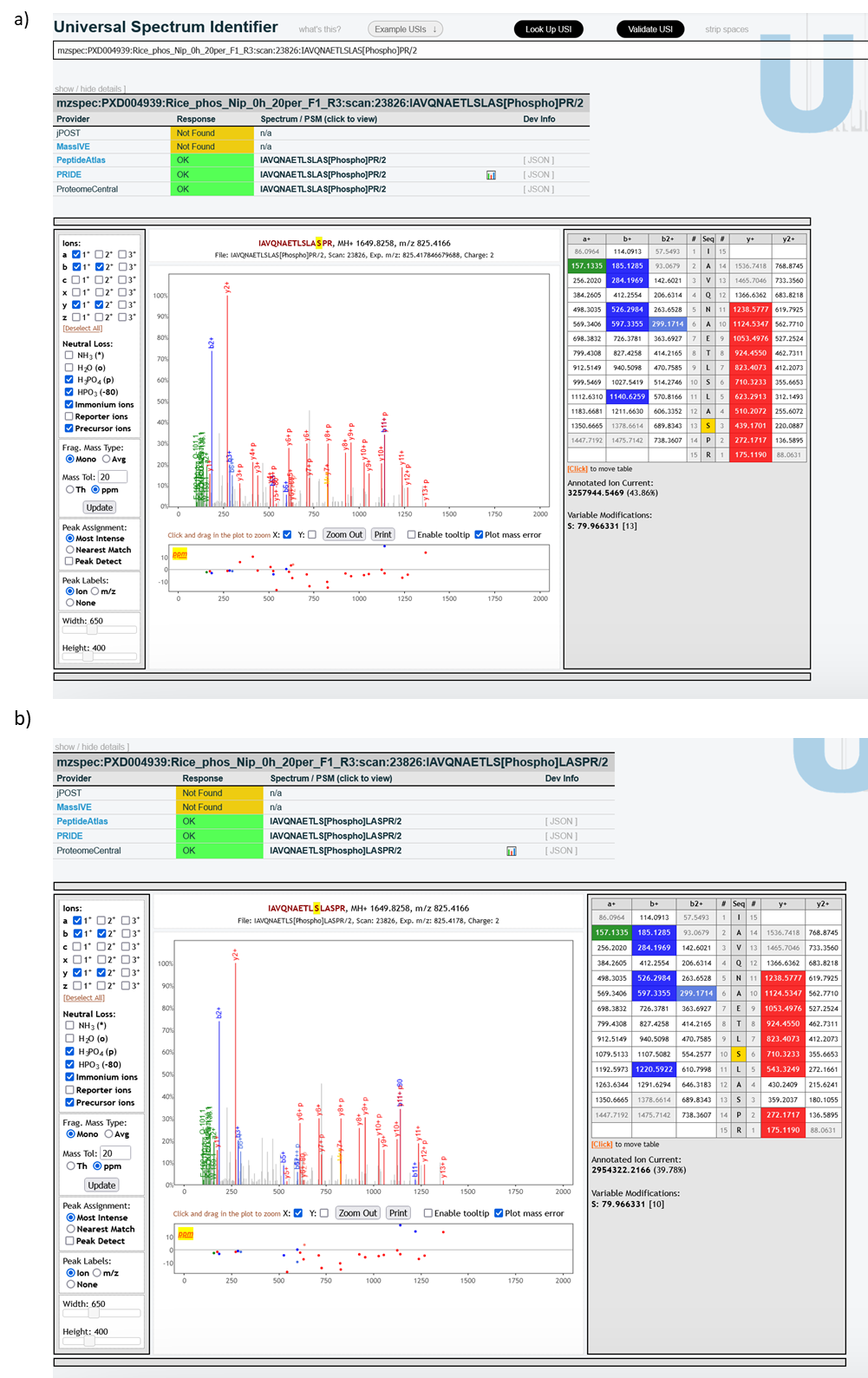

Supplementary Figure 3: Spectrum visualisation comparing the potential phosphorylation positions in the peptide IAVQNAETLSLASPR from the “Rice_phos_Nip_0h_20per_F1_R3” experiment file within the dataset PXD004939. a) shows the spectra with the modification on position 13 (IAVQNAETLSLAS[Phospho]PR), with excellent y-ion coverage supporting the pS site. b) shows the same peptide with modification at position 10 (IAVQNAETLS[Phospho]LASPR), here the spectra has less coverage being seen in the y-ions.

**Motif analysis**

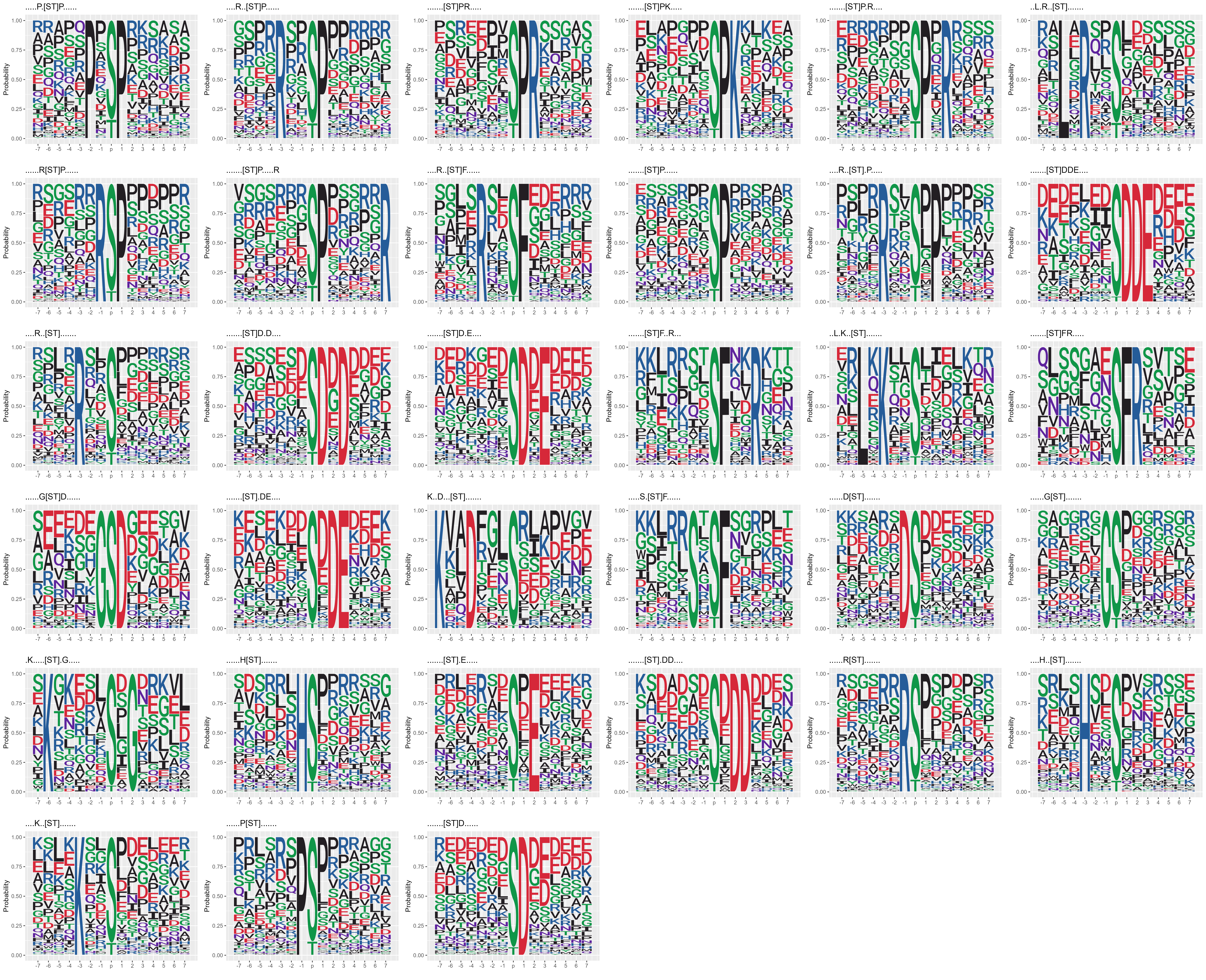

Supplementary Figure 4: Significant motifs surrounding the S/T phosphosites for the gold and silver category sites.

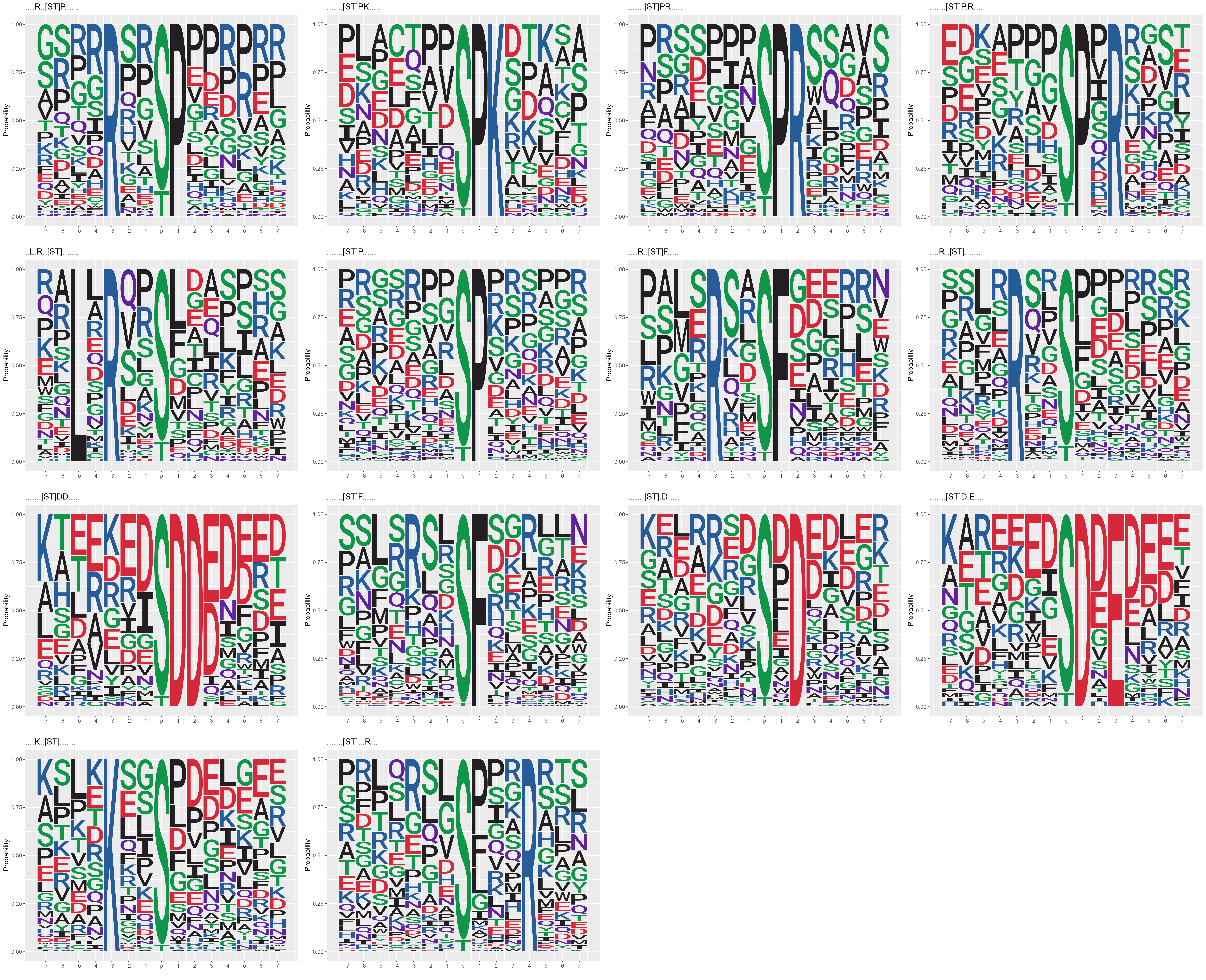

Supplementary Figure 5: Significant motifs surrounding the S/T phosphosites for the gold category sites.

Supplementary Table 2: Table showing the presence/absence of significant motifs in each of the category groupings; Gold, Gold and Silver and Gold, Silver and Bronze.

| Motif | Gold | Gold-Silver | Gold-Silver-Bronze |
| --- | --- | --- | --- |
| ....R..[ST]P...... | 1 | 1 | 1 |
| .......[ST]PK..... | 1 | 1 | 1 |
| .......[ST]PR..... | 1 | 1 | 1 |
| .......[ST]P.R.... | 1 | 1 | 1 |
| ..L.R..[ST]....... | 1 | 1 | 1 |
| .......[ST]P...... | 1 | 1 | 1 |
| ....R..[ST]F...... | 1 | 1 | 1 |
| ....R..[ST]....... | 1 | 1 | 1 |
| .......[ST]DD..... | 1 | 0 | 0 |
| .......[ST]F...... | 1 | 0 | 1 |
| .......[ST].D..... | 1 | 0 | 1 |
| .......[ST]D.E.... | 1 | 1 | 1 |
| ....K..[ST]....... | 1 | 1 | 0 |
| .......[ST]...R... | 1 | 0 | 0 |
| .....P.[ST]P...... | 0 | 1 | 1 |
| ......R[ST]P...... | 0 | 1 | 1 |
| .......[ST]P.....R | 0 | 1 | 0 |
| ....R..[ST].P..... | 0 | 1 | 0 |
| .......[ST]DDE.... | 0 | 1 | 1 |
| .......[ST]D.D.... | 0 | 1 | 1 |
| .......[ST]F..R... | 0 | 1 | 1 |
| ..L.K..[ST]....... | 0 | 1 | 1 |
| .......[ST]FR..... | 0 | 1 | 1 |
| ......G[ST]D...... | 0 | 1 | 0 |
| .......[ST].DE.... | 0 | 1 | 1 |
| K..D...[ST]....... | 0 | 1 | 0 |
| .....S.[ST]F...... | 0 | 1 | 1 |
| ......D[ST]....... | 0 | 1 | 1 |
| ......G[ST]....... | 0 | 1 | 1 |
| .K.....[ST].G..... | 0 | 1 | 0 |
| ......H[ST]....... | 0 | 1 | 1 |
| .......[ST].E..... | 0 | 1 | 1 |
| .......[ST].DD.... | 0 | 1 | 0 |
| ......R[ST]....... | 0 | 1 | 0 |
| ....H..[ST]....... | 0 | 1 | 0 |
| ......P[ST]....... | 0 | 1 | 0 |
| .......[ST]D...... | 0 | 1 | 0 |
| .....P.[ST]P.K.... | 0 | 0 | 1 |
| ....RS.[ST]F...... | 0 | 0 | 1 |
| ......G[ST]P...... | 0 | 0 | 1 |
| ....RS.[ST]....... | 0 | 0 | 1 |
| .......[ST]P...K.. | 0 | 0 | 1 |
| ....R..[ST].D..... | 0 | 0 | 1 |
| ......G[ST]...R... | 0 | 0 | 1 |
| .......[ST]DDD.... | 0 | 0 | 1 |
| ......D[ST]D...... | 0 | 0 | 1 |
| .......[ST].D....K | 0 | 0 | 1 |
| ....H.D[ST]....... | 0 | 0 | 1 |
| .K.N...[ST]....... | 0 | 0 | 1 |
| .......[ST]SP..... | 0 | 0 | 1 |
| .......[ST]E.E.... | 0 | 0 | 1 |
| K....G.[ST]....... | 0 | 0 | 1 |
| ....HS.[ST]....... | 0 | 0 | 1 |
| .....R.[ST]..P.... | 0 | 0 | 1 |
| .......[ST].G..... | 0 | 0 | 1 |
| ......D[ST].E..... | 0 | 0 | 1 |
| .....S.[ST]....... | 0 | 0 | 1 |
| ....HV.[ST]....... | 0 | 0 | 1 |

**Pathway enrichment analysis**

Pathway enrichment analysis was used to identify pathways involved in proteins containing significant motifs identified, against all rice phosphoproteins, using the ClusterProfiler R package. The proteins containing each of the identified enriched motifs, across all categories (Gold, Silver and Bronze), were searched against a background of all phosphoproteins. “RNA binding” is seen commonly enriched across multiple motifs, as is “nuclear speck” and “mRNA splicing, via spliceosome”. Supplementary Figure 6 shows the dotplots of enrichment terms per motif.

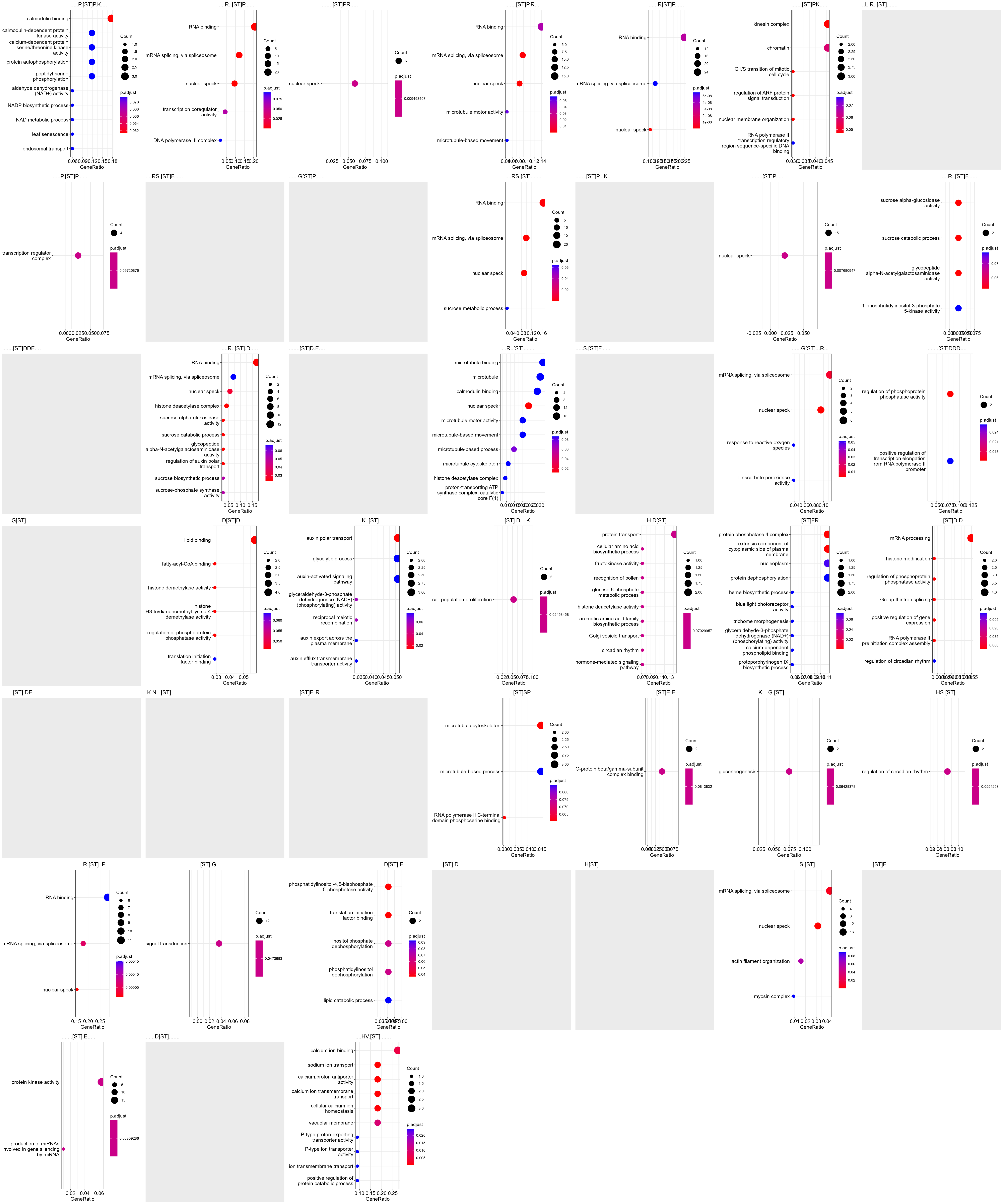

Supplementary Figure 6: Enriched terms for proteins containing each enriched motif across all gold-silver-bronze datasets against all phosphoproteins. Blank plots show motifs which contained no significant motifs.
